# VAPYRIN attenuates defence by repressing PR gene induction and localized lignin accumulation during arbuscular mycorrhizal symbiosis of *Petunia hybrida*

**DOI:** 10.1101/2020.09.16.300590

**Authors:** Min Chen, Sébastien Bruisson, Laure Bapaume, Geoffrey Darbon, Gaëtan Glauser, Martine Schorderet, Didier Reinhardt

**Author notes:** These authors contributed equally (Min Chen, Sébastien Bruisson). Communicating author.

## Abstract

- The intimate association of host and fungus in arbuscular mycorrhizal (AM) symbiosis can potentially trigger induction of host defence mechanisms against the fungus, implying that successful symbiosis requires suppression of defence.
- We addressed this phenomenon by using AM-defective *vapyrin* (*vpy*) mutants in *Petunia hybrida*, including a new allele (*vpy-3*) with a transposon insertion close to the ATG start codon. We explore whether abortion of fungal infection in *vpy* mutants is associated with the induction of defence markers such as cell wall alterations, accumulation of reactive oxygen species (ROS), defence hormones, and induction of pathogenesis-related (PR) genes.
- We show that *vpy* mutants exhibit a strong resistance against intracellular colonization, which is associated with the generation of thick cell wall appositions (papillae) with lignin impregnation at fungal entry sites, while no accumulation of defence hormones, ROS, or callose was observed. Systematic analysis of PR gene expression revealed that several PR genes are induced in mycorrhizal roots of the wild type, and even more in *vpy* plants. Some PR genes are induced exclusively in *vpy* mutants.
- Taken together, these results suggest that *VPY* is involved in avoiding or suppressing the induction of a cellular defence syndrome that involves localized lignin deposition and PR gene induction.

## Introduction

Arbuscular mycorrhiza (AM) is a mutualistic association of the majority of land plants with fungi of the subphylum *Glomeromycotina* (Smith & Read, 2008; Spatafora *et al*., 2016), which confers various benefits to the plant host (Chen, M *et al*., 2018). Although AM require mutual recognition of the partners to establish intracellular compatibility, the interaction is characterized by a very low level of host specificity (Smith & Read, 2008). For example, the AM fungal model species *Rhizophagus irregularis* can colonize most angiosperms with few exceptions represented by plant taxa that are generally incapable of engaging in AM, e.g. the Brassicaceae (including oilseed rape and the model species *Arabidopsis thaliana*) and the Amaranthaceae (represented by spinach and sugar beet).

An striking aspect of the strong compatibility in AM is the massive intracellular colonization of the root cortex by arbuscules and vesicles which can result in >90% colonization of the entire root length of the host. AM fungi share typical fungal cell wall components, e.g. chitin, with fungal pathogens, and plants have very sensitive detection mechanisms for such general microbial molecules, which are known as pathogen-associated molecular patterns (PAMPs), or, more generally, microbe-associated molecular patterns (MAMPs) (Boller & Felix, 2009). Hence, the abundance of fungal material in the root, and the intimate interaction between the two symbiotic partners (Rich *et al*., 2014) imply that disease resistance mechanisms in the host must be under tight control to avoid defence reactions to be triggered against AM fungi (Gianinazzi-Pearson, 1996; Gianinazzi-Pearson *et al*., 1996; Marsh & Schultze, 2001; Zipfel & Oldroyd, 2017).

Several studies have reported induction of defence mechanisms at early stages of AM interactions (Gianinazzi-Pearson *et al*., 1996; Kapulnik *et al*., 1996; Campos-Soriano *et al*., 2010; Marcel *et al*., 2010). After initial induction, these defence responses usually become repressed to, or below, the levels in non-inoculated control roots. Interestingly, in simultaneous inoculations, AM fungi can reduce the induction of defence responses elicited by a pathogen (Guenoune *et al*., 2001), or by a chemical inducer of defence (David *et al*., 1998). This indicates that infection by AM fungi is associated with active suppression of defence. Indeed, an AM fungal effector protein that promotes biotrophic compatibility in host plants by suppressing defence has been identified in *R. irregularis* (Kloppholz *et al*., 2011). On the other hand, it was shown in many cases that the general disease resistance is improved by AMF, also in the aerial parts of colonized plants, a phenomenon known as mycorrhiza-induced resistance (MIR), by a mechanism that resembles induced systemic resistance (ISR) (Jung *et al*., 2012; Pieterse *et al*., 2014). Characteristic markers of defence include the stress hormones salicylic acid (Loake & Grant, 2007), jasmonic acid (Browse, 2009) and ethylene (van Loon *et al*., 2006a), cell wall reinforcements such as callose and lignin (Millet *et al*., 2010; Miedes *et al*., 2014; Chowdhury *et al*., 2016; Liu *et al*., 2018), induction of reactive oxygen species (Jones & Dangl, 2006), and the induction of pathogenesis-related (PR) proteins that are thought to have antimicrobial functions and contribute to disease resistance (van Loon *et al*., 2006b). However, in this context it is important to note that most of our knowledge on plant defence mechanisms were gained in the shoot (mainly from leaves), while defence mechanisms in the roots are considerably different (Chuberre *et al*., 2018), and have been explored to a lesser extent.

We have previously described two allelic mutants in petunia (*Petunia hybrida*), *penetration and arbuscule morphogenesis1-1* (*pam1-1*) and *pam1-2*, which carry transposon insertions in the *VAPYRIN* (*VPY*) gene (further referred to as *vpy-1* and *vpy-2*, respectively). *Vpy-1* and *vpy-2* mutants are defective in intracellular accommodation of AM fungi (Sekhara Reddy *et al*., 2007; Feddermann *et al*., 2010). Based on detailed phenotypic analysis, VPY is involved in accommodation of AM fungi at the two intracellular stages, during infection of epidermal and hypodermal cells, and during arbuscule formation. VPY function is conserved between petunia and *Medicago truncatula* (Pumplin *et al*., 2010), and acts downstream of calcium spiking, the central element in symbiotic signaling (Murray *et al*., 2011). Interestingly, VPY protein is localized to small mobile compartments, which are thought to be involved in cellular trafficking during symbiosis (Feddermann *et al*., 2010; Pumplin *et al*., 2010; Zhang *et al*., 2015; Bapaume *et al*., 2019; Liu *et al*., 2019). The fact that the AM fungus in *vpy* mutants exhibits conspicuous deformations upon cell penetration, and forms hyphal septa (a sign of stress) (Sekhara Reddy *et al*., 2007; Feddermann *et al*., 2010), suggest that this trafficking pathway may be involved, directly or indirectly, in modulating defence during intracellular stages of AM.

Here, we describe a new mutant allele of the petunia *VPY* gene, *vpy-3*, which is a null allele, due to a *dTph1* transposon insertion after only eight codons from the start codon. *Vpy-3* exhibits similar defects as the two other alleles *vpy-1* and *vpy-2*, indicating that these also represent functional null alleles. Microscopic and molecular analysis using a range of defence markers indicates that the abortion of the AM fungus in *vpy* mutants involves a cellular defence response that is independent of callose deposition and of the classical stress hormones salicylic acid, jasmonic acid and ethylene, but is associated with induction of several PR genes, and with cell wall lignification during intracellular invasion.

## Materials and Methods

### Plant lines, fungal material, and growth conditions

Seeds of the petunia transposon line W138 were germinated on seedling substrate (Klasmann, http://www.klasmann-deilmann.com). After four weeks, plantlets were transferred to a sterilized mixture of 75% sand with 25% unfertilized soil (further referred to as sand substrate), inoculated with around 10g of pot culture inoculum of *Rhizophagus irregularis* (MUCL 43204), and cultured as described (Nouri *et al*., 2014). Nurse-plant inoculation was carried out by co-culturing in the same pot petunia mutant plants with chive plants (*Allium schoenoprasum*) that had been inoculated at least four weeks before. Plants were grown in growth chambers with a day/night cycle of 12h (25°C)/12h (20°C).

### Isolation of the *vpy-3* mutant allele

The *vpy-3* allele was isolated from the transposon line W138 of *Petunia hybrida* as described (Sekhara Reddy *et al*., 2007; Rich *et al*., 2015). Briefly, eight individuals per segregating family were assessed for mycorrhizal colonization after 5 weeks of colonization with *R. irregularis* (primary screen). Root samples were taken from inoculated plants, stained with trypan blue and screened visually for the presence of AM fungal structures. Families with AM-defective individuals were further grown for seed production and additional seeds of the respective family were sown for phenotypic analysis and assessment of the segregation pattern (secondary screen).

Homozygous mutant individuals of the new mutant line were crossed with *pam1-1* and *pam1-2* (Feddermann *et al*., 2010) to test for allelism. All progeny of these crosses exhibited a similar mutant phenotype as the parents, indicating that the new mutant carries a mutation in the *PAM1* gene. Since the gene encodes the VAPYRIN protein, we further refer to the mutant as *vpy-3*. Isolation of the *vpy-3* locus by PCR, cloning into pGEMT, and sequencing revealed an insertion of a *dTph1* copy after 25 nucleotides from the start codon (ATG). For detailed phenotypic analysis, the *vpy-3* allele was stabilized by segregating out the active translocator locus *ACT1* after crossing with the stabilizer line W5 (Stuurman & Kuhlemeier, 2005).

### Assessing AM fungal colonization and papilla formation

For primary mutant screening, roots were harvested, washed and directly stained with Trypan Blue (0.01% w/v) in 0.5% (v/v) acetic acid for 10 min at 95°C, and washed with water for visual inspection. For the secondary screening, roots were cleared in 10% KOH (30 min at 95°C), washed twice with water, stained for 10 min with Trypan Blue staining solution at 95°C (20% glycerol, 30% lactic acid and 0.01% Trypan Blue) and rinsed twice with 10% lactic acid.

For initial assessment of fungal hyphae and cell wall papillae in *vpy-3* (Fig. 2), plants were inoculated with *R. irregularis* in nurse plant chambers for four weeks. Roots were fixed for 2h at room temperature in 4% (v/v) paraformaldehyde. After several washes, roots were stained overnight at 4°C in the dark with 5 µg/ml wheat germ agglutinin (WGA) coupled to FITC (www.lifetechnologies.com) in Soerensen’s phosphate buffer (0.133 M, pH=7.2). Before mounting, samples were incubated for 10 min in 50 µg/ml propidium iodide at room temperature. Images were acquired on a Leica SP5 confocal microscope. In a second experiment for quantification of papilla formation in all *vpy* alleles, plants were inoculated with *R. irregularis* in nurse plant chambers for four weeks, then roots were harvested and cleared in 10% KOH for 20 min. After four washes with deionized water, roots were stained overnight with WGA-Alexa488 in Soerensen’s phosphate buffer (0.133 M, pH=7.2), followed by counterstaining with 0.2% basic fuchsin (Sigma, 857343). For microscopy, the roots were immersed in a modified version of ClearSee (Kurihara *et al*., 2015) containing 10% (w/v) xylitol, 25% (w/v) urea, and 2% (w/v) SDS.

**Figure 1.**
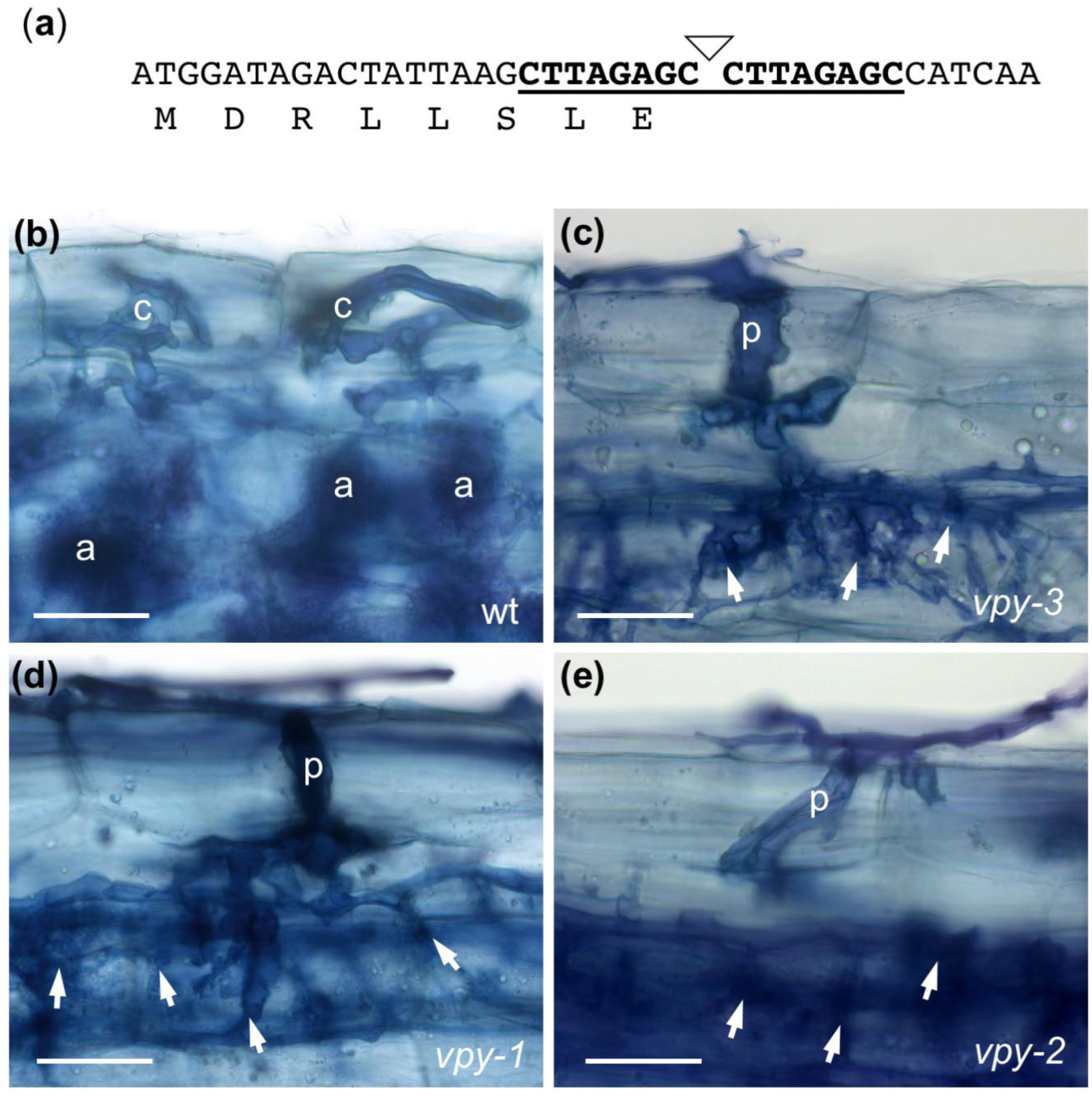
AM-defective phenotype and molecular characterization of *vpy-3*. (a) Molecular aspects of the new *vpy-3* allele. The petunia *VAPYRIN* coding sequence is shown from the ATG start codon to the insertion site of a *dTPh1* transposon (triangle). The 8 bp target site duplication is bold and underlined. Due to stop codons in all three reading frames of the *dTPh1* sequence, the truncated protein can be predicted to contain only 8 residual amino acids of VAPYRIN. (b) Colonization pattern in the wild type revealed by Trypan-Blue staining. Infection by hyphal coils (c) in hypodermal cells, and arbuscules (a) in the cortex. (c-e) Colonization pattern in *vpy* mutants revealed by Trypan-Blue staining. Infection by enlarged penetration peg (p), and subsequent profusely branched hyphal colonization (asterisks) without arbuscules in *vpy-3* (c), and for comparison, *vpy-1* (d), and *vpy-2* (e). Scale bars: 50 μm

**Figure 2.**
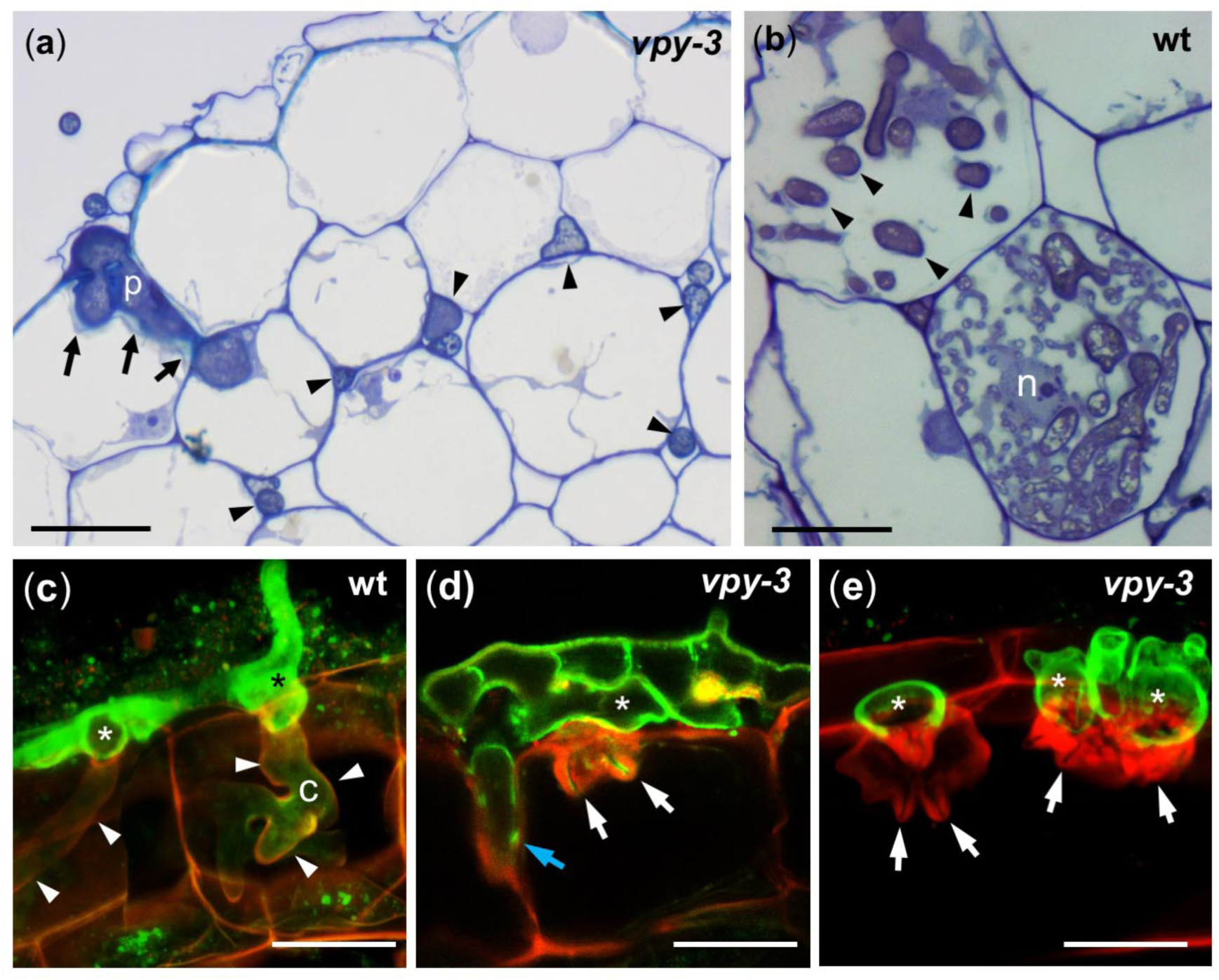
Cell wall appositions in *vpy-3* mutants infected with *R. irregularis*. (a) Colonization pattern in *vpy-3* mutant revealed by semithin sections of resin-embedded material stained with Toluidine Blue. A hyphopodium with an intercellular infection peg (asterisk), and cell wall appositions (arrows). Fungal colonization is mostly restricted to the intercellular spaces (arrowheads). (b) Colonization of the wild type. Cortical cells contain abundant intracellular hyphae (arrowheads), and finely branched arbuscules, with the host nucleus (n) at a central position. (c-e) Confocal scanning micrographs of inoculated root hypodermal cells with the fungus stained by fluorescein isothiocyanate-labeled wheat germ-agglutinine (FITC-WGA, green), and the plant cell walls by propidium iodide (PI, red). (c) In the wild type, two fungal hyphopodia (asterisks) successfully penetrated two adjacent hypodermal cells. The infection hyphae are surrounded by a thin layer of plant extracellular matrix material (arrowheads). (d) On a *vpy-3* mutant plant, the fungus has formed a highly septate complex hyphopodium (asterisk) from which it has attempted to penetrate a hypodermal cell. Thick cell wall appositions surround the fungal penetration hyphae (white arrows). A penetration hypha has inserted itself between two adjacent cells (blue arrow) (e) Extreme case of an inoculated *vpy-3* mutant as in (d) with massive cell wall appositions that have completely blocked fungal penetration. Scale bars: 40 μm

### Callose and lignin staining

Wild type and *vpy* mutants were inoculated with nurse plants for four weeks. For callose detection, roots were fixed overnight with 4% (v/v) paraformaldehyde (**Fig. 6**), or with 1:3 acetic acid:ethanol (**Table S1**), washed 5x with water, followed by staining with 0.01% (w/v) aniline blue in 150 mM KH_2_PO_4_ pH=9.5 for 48h. Epifluorescence images were acquired with a Leica DMR microscope equipped with an Axiocam (Zeiss). Lignin was stained with phloroglucinol solution (100 ml 95% EtOH, 16 ml HCl conc., 0.1g phloroglucinol) for 30 min and directly analyzed.

### Detection of reactive oxygen species (ROS)

Plants were inoculated with nurse plants for 4 weeks. For detection of H_2_O_2_, roots were harvested and treated with a freshly prepared solution of 1mg/ml 3,3’-Diaminobenzidine (DAB) in 10 mM MES-NaOH (pH 5.6) (Salzer *et al*., 1999). Preparing the solution requires intense stirring and careful acidification with HCl to ca. pH=3 (to increase DAB solubility) followed by buffering with MES-NaOH. After overnight incubation at room temperature in the dark, roots were washed 4x with water and incubated in ClearSeeD for microscopic analysis. For detection of O_2_^--^, roots were treated for 1h at room temperature in the dark with a fresh solution of 1mg/ml Nitroblue tetrazolium (NBT) in 10 mM phosphate buffer, pH=7.8. After four washes with water, roots were incubated in ClearSeeD for microscopic analysis.

### Electron microscopy and preparation of semithin sections

Wild type and *vpy-3* mutants were inoculated with *R. irregularis* in nurse plant chambers for 4 weeks. Roots were fixed for 2h at room temperature in 4% (v/v) glutaraldehyde and postfixed with 1% (w/v) OsO_4_ at 4°C overnight. Further processing of the samples and embedding in Spurr’s resin was carried out as described (Spurr, 1969). Semithin sections (1 µm) were stained with 1 % (w/v) toluidine blue in 1% (w/v) borax (Na_2_B_4_O_7_). Images were acquired in the bright field mode on a Leica DMR microscope equipped with an Axiocam (Zeiss). For transmission electron microscopic (TEM) analysis, ultrathin sections (70 nm) were prepared on a Reichert-Jung Ultracut E. Contrasting was performed with 2% (w/v) uranyl acetate (UO_2_(CH_3_COO)_2_) and lead citrate solution prepared according to (Reynolds, 1963). Images were acquired on a Philips Biotwin CM100.

### β-1,3-Glucanase immunostaining

Wild type an *vpy-3* plants were inoculated with nurse plant inoculum of *R. irregularis*. Root material was fixed for 2h with 2% (w/v) paraformaldehyde and 1% (v/v) glutaraldehyde at room temperature. Fixed material was dehydrated and embedded in Lowicryl K4M (www.sigmaaldrich.com) as described (Altman *et al*., 1984) with the following modifications: embedding involved a series of lowicryl diluted with 95% (v/v) ethanol as follows: 1:2 (vol:vol) for 2h, 1:1 for 2h, 2:1 overnight, followed by twice 100% lowicryl for 2h and polymerization at 55°C. Ultrathin sections (70 nm) were blocked with 20 mM NH_4_Cl, 20 mM lysine and 20 mM glycine. Rabbit antiserum raised against extracellular glucanase (Beffa *et al*., 1993) was used at a dilution of 1:5 in blocking agent. Goat-anti-rabbit antibodies coupled to 10 nm gold beads (http://www.bbisolutions.com) served as secondary antibodies. Contrasting was performed with 2% (w/v) uranyl acetate (UO_2_(CH_3_COO)_2_) and lead citrate solution prepared according to (Reynolds, 1963). Controls without the primary antibody did not show any labeling.

For quantification of immunogold signal, for each treatment six representative pictures of hypodermal cells from two independent experiments were printed out at the same magnification. A 2D grid of small crosses at regular distances (1 cm) printed on a transparent foil was overlaid and the gold particles were counted in the respective cellular compartments relative to the number of crosses within the same compartment.

### Identification of PR genes and lignin biosynthetic genes

PR gene candidates for quantitative real-time reverse-transcriptase PCR (qPCR) were identified by searching the *Petunia axillaris* predicted transcriptome at the SolGenomics database (https://solgenomics.net) using tobacco PR proteins (van Loon *et al*., 2006b) by tblastn. Primers were designed for all close homologues for a preliminary analysis of gene expression in mycorrhizal and non-mycorrhizal roots of wild type and *vpy* mutant roots to identify genes that were expressed in any of the tested conditions, and these were used for qPCR analysis. To identify lignin biosynthetic genes with a potential role in roots, we identified all genes annotated as lignin-related in a previous EST (expressed sequence tag) sequencing project (Breuillin *et al*., 2010). The full-length gene sequences were than identified from the predicted transcriptome of P. axillaris, and used for primer design. All genes were analyzed in mycorrhizal and non-mycorrhizal wild type and mutant roots, and genes that were expressed at any of these conditions were further assessed by qPCR.

### RNA extraction and quantitative real-time RT-PCR

For gene expression analysis, plants were either inoculated for 4 weeks with nurse plants (**Tables 1**,**2**), or wild type plants were treated with chitin oligosaccharides or with a *Penicillium* preparation (Thuerig *et al*., 2005) for 1 and 4h at concentrations of 1 μg/ml and 10 μg/ml, respectively (**Table 3**). Frozen petunia roots were placed in 2 mL Eppendorf tubes containing a glass bead and were ground using a ball mill. Total RNA was extracted from the powdered roots according to the protocol of the Direct-zol RNA miniprep kit from Zymo Research (https://www.zymoresearch.com), using trizol solution for lysis (38% v/v), saturated phenol (pH 8), 0.8 M guanidine thiocyanate, 0.4 M ammonium thiocyanate, 0.1M Na-acetate pH 5, and 5% (v/v) glycerol. The Direct-zol RNA kit involves a DNAse step to remove genomic DNA during RNA extraction. RNA was used for reverse transcription according to the protocol of the SensiFASTtm cDNA synthesis kit from Bioline. PCR reactions were carried out with 5 μL of 100x diluted cDNA solution, 1 μL of 10 μM forward and reverse primer, 12.5 μL of SensiMix SYBR Hi-ROX (BIO-RAD) and DNAse/RNAse free water up to 15 μL. The reaction cycle was 95°C for 10 min followed by 45 amplification cycles (95°C for 20s, 64°C for 20s, 72°C for 20s). All samples were analyzed in technical duplicates from eight independent replicate plants. Actin and glyceraldehyde-3-phosphate dehydrogenase (GAPDH) were used as reference genes. Relative expression values were calculated using the delta-Ct method (Pfaffl, 2001), and log_2_ transformed for further analysis. Data are expressed as -fold changes in relative gene expression (AM/control) in Table 1. All data were statistically analyzed using two-way ANOVA and Tukey’s HSD test with p-value < 0.05.

**Table 1.**
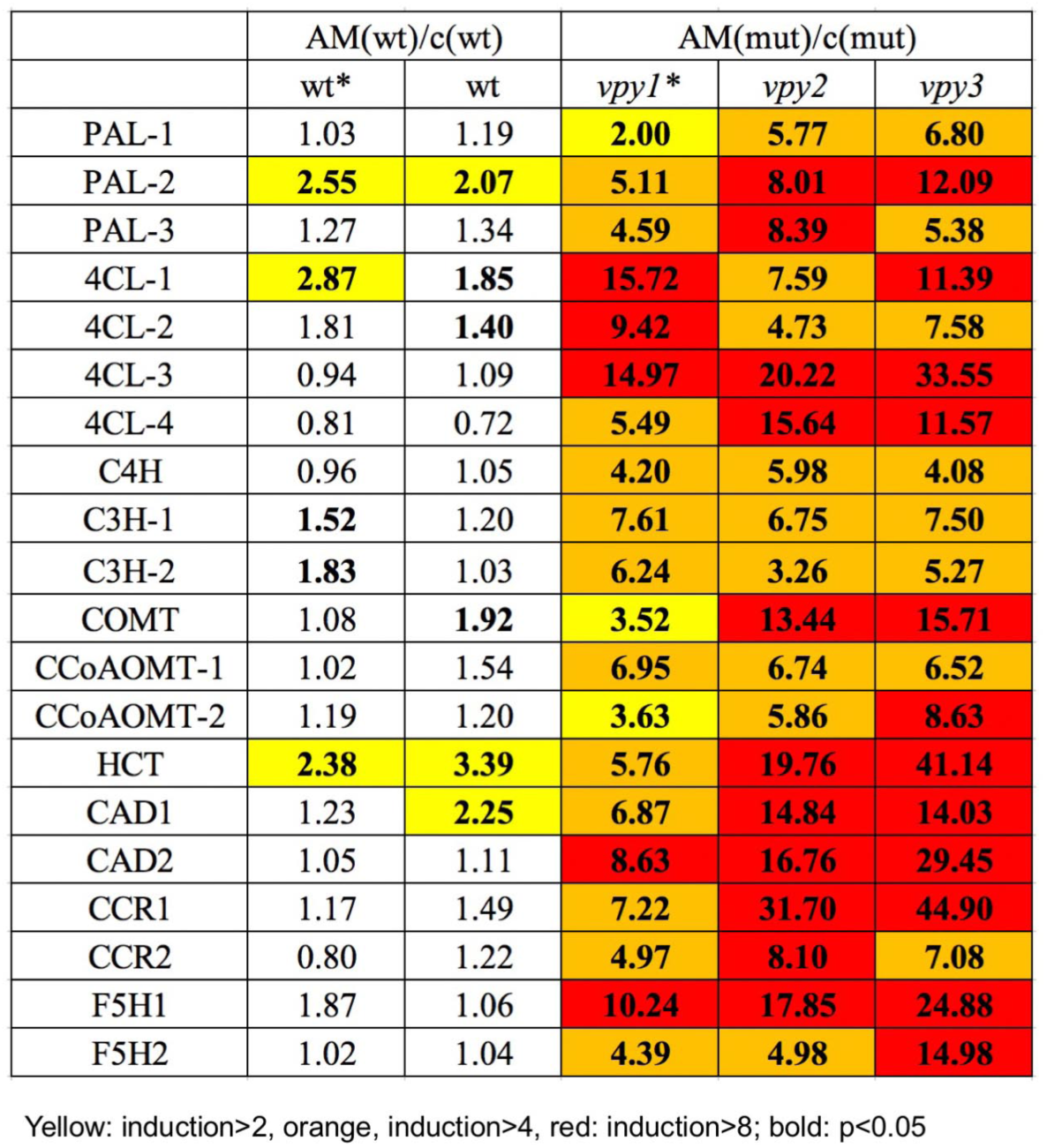
Expression ratios of lignin-related genes in wild type and *vpy* mutants relative to the respective non-mycorrhizal controls. Induction of lignin biosynthetic genes was determined by qPCR with actin and glyceraldehyde-3-phosphate dehydrogenase (GAPDH) as reference genes. Values represent -fold induction ratios derived by dividing the expression values of mycorrhizal plants by values of non-mycorrhizal controls (both normalized with the two reference genes) in wt, *vpy-1, vpy-2*, and *vpy-3*. All expression values were derived from 8 biological replicates. Color shading represents induction >2-fold (yellow), >4-fold (orange), and >8-fold (red). Significant induction ratios are indicated by bold type font (two-way ANOVA).

### Determination of salicylic acid and jasmonic acid

Levels of JA and Ile-JA were determined by ultra-high performance liquid chromatography-tandem mass spectrometry (UHPLC-MS/MS) according to {Glauser, 2014 #5609}. SA and conjugated SA were separated by two-phase extraction, followed by acid hydrolysis of conjugated SA, and quantification as described {Fragniere, 2011 #5675}.

## Results

### Isolation of a new *pam1*/*vpy* allele

In a forward genetic screen for AM-defective mutants, we isolated a mutant candidate with severely decreased colonization levels of the AM fungus *Rhizophagus irregularis* (<5% total root length colonized). Colonization of the root surface resulted in the production of deformed hyphopodia and penetration of the epidermis and hypodermis was often arrested. Based on the phenotypic similarity with the previously isolated mutants *pam1-1* and *pam1-2* (Sekhara Reddy *et al*., 2007; Feddermann *et al*., 2010), we crossed the new mutant with these *pam1* alleles to test for allelism. 100% of the F1 progeny showed the same AM-resistant phenotype as the two parents (data not shown), confirming that the new mutant is allelic to *pam1-1* and *pam1-2*. Since the encoded protein has since been named VAPYRIN (VPY) due to its domain structure (Feddermann *et al*., 2010; Pumplin *et al*., 2010; Feddermann & Reinhardt, 2011; Murray *et al*., 2011), we further refer to the new allele as *vapyrin-3* (*vpy-3*), and the previously isolated alleles were renamed to *vpy-1* and *vpy-2*, respectively.

The coding region of the *VPY* gene in the *vpy-3* mutant was amplified by PCR and cloned in order to identify the nature of the mutation. Indeed, a *dTph1* insertion was detected at position 25 from the predicted start codon (ATG), leaving only 8 of the 535 amino acids of the predicted protein (Feddermann *et al*., 2010) (**Fig. 1a**). The *dTph1* sequence contains multiple stop codons in all three reading frames in both orientations, hence, read-through over the transposon sequence is unlikely, and therefore, *vpy-3* likely represents a null allele.

Initial phenotypic analysis of *vpy-3* mutants, alongside with *vpy-1* and *vpy-2*, showed defects at early stages of infection after inoculation from nurse plants (nearby growing inoculated wild type plants). While in the wild type, *R. irregularis* forms hyphal coils in hypodermal cells (Rich *et al*., 2014) (**Fig. 1b**), all three mutants exhibited thickened infection pegs that seemed to penetrate the hypodermis directly, instead of forming a hyphal coil (**Fig. 1c-e**). Subsequent colonization of the cortex resulted in the formation of arbuscules in the wild type (**Fig. 1b**), whereas all three mutants showed similar unstructured profusely growing hyphal material **(Fig. 1c-e**, arrows). However, all three mutant genotypes reached similar colonization levels as the wild type, after inoculation with nurse plants (>90% total root length colonization) {Sekhara Reddy, 2007 #2695}.

### *vpy* mutants exhibit cell wall alterations at sites of mycorrhizal infection

In order to distinguish with confidence intra- and intercellular fungal structures in *vpy-3*, semi-thin sections of resin-embedded material were prepared. They revealed that fungal hyphae in *vpy-3* were mostly confined to the apoplast at all stages of infection from the rhizodermis to the cortex (**Fig. 2a**, arrowheads), whereas the wild type showed extensive intracellular colonization by hyphal coils and arbuscules (**Fig. 2b**). Interestingly, hypodermal cells adjacent to fungal penetration hyphae exhibited thickened cell walls in *vpy-3* (**Fig. 2a**, arrows).

To better distinguish the plant and fungal structures, and to clarify their relationships during infection, we performed confocal microscopic analysis with a double staining protocol that allows to differentially stain plant and fungal cell walls based on their biochemical constituents. Wild type and mutants plants inoculated from nurse plants were stained with propidium iodide (PI) for plant cell walls, and with wheat germ agglutinine coupled to fluoresceinisothiocyanate (WGA-FITC) which stains fungal structures (**Fig. 2c-e**). In wild type plants, the fungus formed hyphopodia on the root surface, from which it invaded hypodermal cells (**Fig. 2c**). Intracellular hyphae were surrounded by a thin layer of interfacial material (**Fig. 2c**, arrowheads). In contrast, hyphopodia on *vpy-3* mutants were enlarged and septate (**Fig. 2d**,**e**). When fungal infection hyphae in the mutant produced projections into hypodermal cells, they remained short and were surrounded by thick cell wall appositions from the host (**Fig. 2d**,**e**; arrows). Occasionally, penetration proceeded between hypodermal cells in the apoplastic space (**Fig. 2d**, blue arrow, compare with Fig. 2a).

In order to quantitatify the cellular defence response we performed a second experiment with all three *vpy* alleles, using a slightly modified staining protocol involving basic fuchsin staining for plant cell walls, and WGA-ALEXA488 for fungal structures. Again, prominent cell wall appositions were observed when mutants were infected by *R. irregularis*. Formation of local cell wall appositions that resembled papillae (Chowdhury *et al*., 2014) was particularly strong in hypodermal cells of the mutants (**Fig. S1b-d**), whereas the wild type hypodermal cells allowed invasion and growth of infection hyphae without restriction (**Fig. S1a**). All three mutant alleles exhibited similar patterns of cell wall alterations, and quantification showed that the relative number of papillae was similar in all three *vpy* alleles, whereas papillae were almost never observed in the wild type (**Fig. S1e**).

### Fungal passage through the hypodermis is blocked in *vpy-3*

To assess in more detail the cellular aspects of the early stages of infection, we performed transmission electron microscopy (TEM) on mycorrhizal wild type and *vpy-3*. First, the stages of early infection were assessed in the wild type. After random proliferation on the root surface, *R. irregularis* hyphae grew along the furrows between adjacent epidermal cells (**Fig. 3a**). After this initial contact stage, the hyphae produced thick-walled swellings in the space between adjacent epidermal cells, often in contact with the external surface of a subtending hypodermal cell (**Fig. 3b**). These represent the hyphopodia, from which the hypodermal cells were invaded, This process involved the breaching of external layers of the wall (**Fig. 3c**, white arrows), while the interior layers of the cell wall extended along the infection hypha to produce a continuous apoplastic sleeve that contained the fungal infection hypha (**Fig. 3c**, black arrows), (compare with (Rich *et al*., 2014)). On *vpy-3* mutant roots, *R. irregularis* also attempted to insert at the clefts between adjacent epidermal cells, and produced thick-walled structures that resembled hyphopodia, however, they appeared more irregular than hyphopodia in the wild type, consistent with the previously observed fungal structures revealed by Trypan Blue and WGA-Alexa488 (**Fig. 1**,**2**). After forcing their way to the surface of the hypodermal cells, fungal hyphae penetrated the hypodermal cell wall in a fashion that superficially resembled the invasion of wild-type cells (compare **Fig. 3c** and **Fig. 3e**), since in both cases, the outer layer of the cell wall (slightly more electron-translucent) was breached, and the hyphae enetered the cellular lumen. However, in contrast to the wild type, the mutant roots produced thick wall appositions that surrounded the inserting hyphae (**Fig. 3e**, arrowheads). Subsequent stages of cell invasion appeared to face resistance, since the fungal hyphae grew rather irregularly, instead of forming a typical hyphal coil (**Fig. 3f**). Progressive deposition of electron-dense cell wall material around the fungus continued (**Fig. 3f, arrowheads**), and occasionally resulted in extreme cell wall masses (**Fig. 3g, arrowheads**), while the fungal structures appeared collapsed and devoid of cytoplasm, indicating that the fungus was dead (**Fig. 3g, asterisks**). In general, hypodermal cells have a considerably thicker cell wall than epidermal and cortical cells (**Fig. 3**) (Rich *et al*., 2014) indicating that the hypodermis may represent a barrier for fungal access to the cortex. Also, epidermal cells often showed signs of degeneration and collapse in both genotypes, indicating that in aging roots, the hypodermis functionally replaces the epidermis and takes over its protective role as the outermost cell layer. Taken together, these results and the confocal analysis (**Fig. 2; Fig. S1**) indicate that in *vpy* mutants, the AM fungus meets a pronounced local defence response at the site of attempted root penetration.

**Figure 3.**
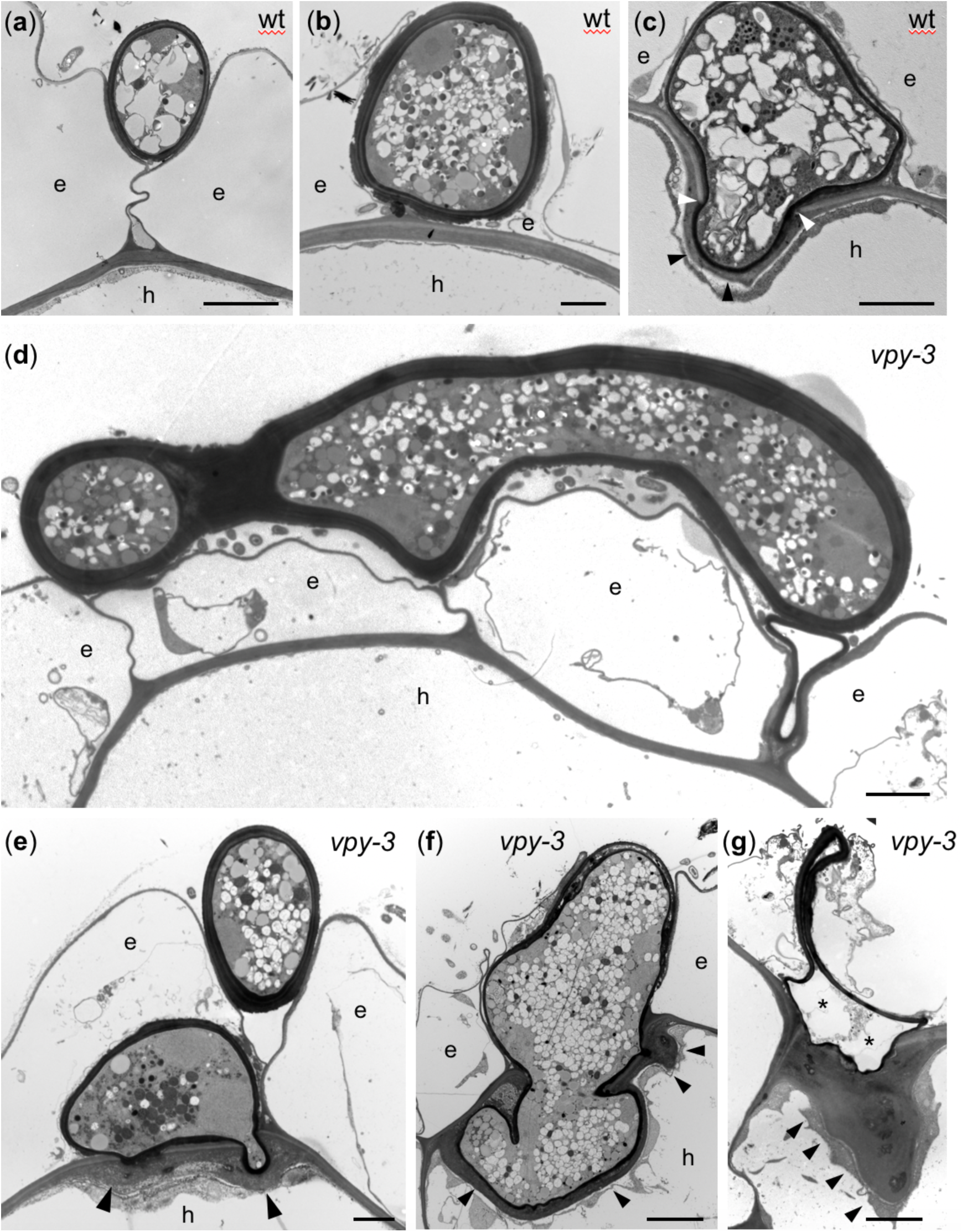
Ultrastructural analysis of root penetration in wt and *vpy-3*. Transmission electron micrographs of transverse sections of wildtype (a-c), and *vpy-3* mutants (d,e) inoculated with *R. irregularis*. All fungal structures are characterized by an electron-dense cell wall that appears almost black. (a) Fungal hypha growing in the cleft between adjacent epidermal root cells. (b) A thick-walled hyphopodium formed above a hypodermal cell (h) between two adjacent epidermal cells. (c) Hyphal entry point into a hypodermal cell (h). Electron translucent layers of the original cell wall were breached be the AM fungus (white arrowheads). The infection hypha is surrounded by a plant-derived layer of cell wall matrix material (black arrowheads). (d) Fungal hypha growing on the root surface of *vpy-3* and attempting to insert between adjacent epidermal cells. (e) Extraradical hypha inserted between two adjacent epidermal cellsof *vpy-3* (top), and hyphopodium-like structure infecting a subtending hypodermal cell (h). At the entry points, the host has deposited thick layers of cell wall material (arrowheads). (f) Attempted entry into a hypodermal cell results in a distorted hyphal clump that is surrounded by an electron-dense layer of cell wall with thickened regions (arrowheads). (g) Aborted entry in a *vpy-3* mutant with an extremely thick cell wall papilla at the site of attempted penetration (arrowheads). The fungal hyphopodium (asterisks) has collapsed and has lost most of its content. Scale bars, 2 μm.

### AM fungal penetration triggers papilla-like cell wall appositions in the cortex of *vpy-3*

Since *vpy* mutants have a distinct arbuscule phenotype (**Fig. 1c-e, Fig. 2a**) (Sekhara Reddy *et al*., 2007), we assessed the AM fungal colonization pattern in the cortex of *vpy-3* in more detail by TEM analysis. In wild type plants, penetration hyphae (PH) entered cortical cells by breaching the cell wall as in the case of hypodermal cells (**Fig. S2a**, white arrowheads), and a thin layer of interfacial cell wall material from the host was deposited on the fungal cell wall (**Fig. S2a**, black arrowheads). Intracellular structures of arbuscules, such as trunk hypha (TH) and fine branches (asterisks) were surrounded by a peri-arbuscular membrane, but hardly any interfacial cell wall material was observed between the membrane and the fungal cell wall (**Fig. S2a, b**). Interestingly, this was also the case for cells that contained collapsing abuscule branches (**Fig. S2c**), indicating that the degenerating fungus did not cause the deposition of host cell wall material around the fungal corpses, although it might be expected to release large amounts of fungal MAMPs that could potentially trigger a defense response. In *vpy-3* mutants, in contrast, attempted cell penetration elicited the formation of local papillae that prevented hyphal entry (**Fig. S2d**). Confocal microscopy confirmed that *vpy-3* mutants formed papillae around fungal structures, in contrast to wild type cells with arbuscules (**Fig. S2e**,**f**).

### Infection of *vpy* mutants by R. irregularis does not trigger accumulation of reactive oxygen species

A widely observed symptom of pathogen infection is the formation of reactive oxygen species (ROS), a reaction known as oxidative burst (Torres *et al*., 2006). We used 3,3’-diaminobenzidine (DAB) staining to visualize hydrogen peroxide (H_2_O_2_) accumulation during AM development (Daudi & O’Brien, 2016). DAB staining revealed a general weak background in both, wild type and *vpy* mutant roots (**Fig. S3**). Since confocal and TEM analysis had revealed the strongest defence reactions in hypodermal cells, we focused our attention on fungal penetration sites in this cell type. Wild type hypodermal cells with an infection hypha showed strongest DAB signal in the hyphae, rather than in the host cytoplasm (**Fig. 4a**). Similarly, penetrated *vpy* mutants exhibited strongest DAB staining inside the fungal hyphae (**Fig. 4b-d**). Notably, cell wall appositions of the mutant host did not exhibit elevated signal (**Fig. 4b-d**, arrowheads), suggesting that the cellular defence response in *vpy* mutants does not involve H_2_O_2_ accumulation. Further examination of cortical colonization revealed that also at later stages of AM development, strongest DAB signal was associated with fungal structures (**Fig. S3**). In the case of arbuscules in the wild type, it was impossible to distinguish whether the signal was localized to the fungus or the surrounding host cytoplasm (**Figure S3d**,**e**), however, in the case of vesicles, the signal was clearly confined to the fungal cytoplasm (**Fig. S3c**,**f**). While the staining was comparable in vesicles of *vpy-2* and wild type roots (compare **Figs. S3c and S3f**), it was consistently weaker in cells with abnormal arbuscules in mutants, vs. fully developed arbuscules in the wild type (compare **Figs. S3b and S3d**). This staining pattern was consistent in all three *vpy* alleles.

**Figure 4.**
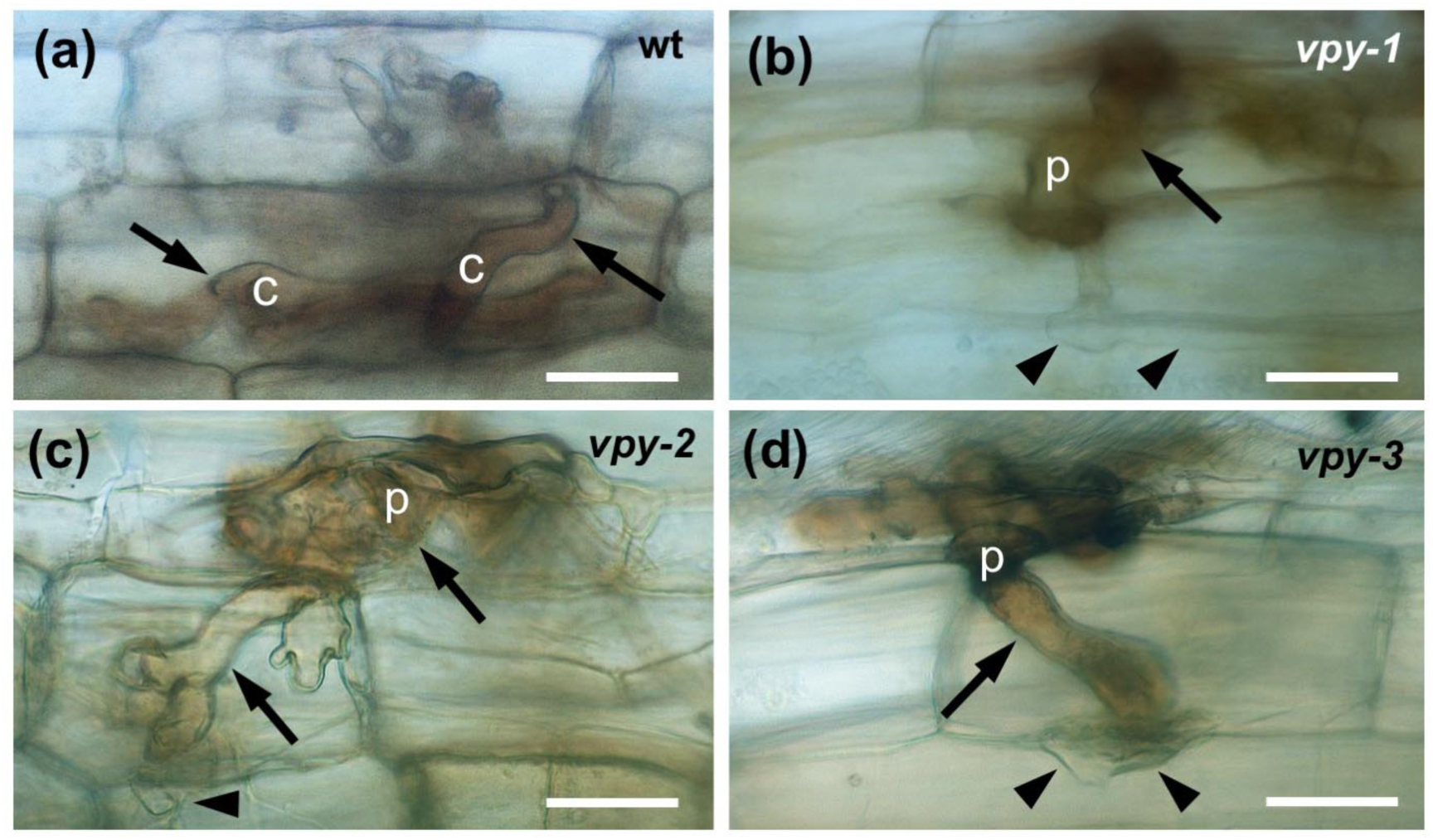
Cytochemical detection of H_2_O_2_ during infection of *vpy* mutants. Plants were infected from mycorrhizal nurse plants, and entry points in hypodermal cells of wild type (a), *vpy-1* (b), *vpy-2* (c), and *vpy-3* (d), were evaluated after staining with diaminobenzidine (DAB). Strongest staining was observed in the fungal cytoplasm (arrows), whereas host cell walls and cell wall appositions (arrowheads) exhibited only weak background signal. Size bar 25 µm.

Next, we performed nitroblue tetrazolium (NBT) staining (Kumar *et al*., 2014) to evaluate whether AM colonization is associated with O_2_^-^ accumulation. In general, NBT staining produced very strong signals in and around the root meristem in wild type as well as in *vpy* mutant roots, irrespective of their mycorrhizal status (**Fig. S4a-e**). Considering infection sites in colonized wild type roots, NBT staining was essentially restricted to fungal structures (**Fig. S4f**). Similarly, infected hypodermal cells in *vpy-1* mutants showed NBT staining almost exclusively in fungal hyphae (**Fig. S4g**), and the same was true for fungal colonization in the cortex (**Fig. S4h**). Corresponding infection sites in *vpy-2* and *vpy-3* mutants showed very similar staining pattern as in *vpy-1*, with staining only in fungal structures (**Fig. S4i**,**j**). Taken together, it appears that ROS levels are higher in the fungal cytoplasm than in the host (except for the root tips), and that infected cells of *vpy* mutants did not accumulate higher levels of ROS than the corresponding wild type cells.

### Abortion of fungal entry in *vpy* mutants does not correlate with callose deposition

We next tested whether papillae in *vpy* mutants contain callose, a β-1→3-linked glucose polymer, that is often associated with cellular defense (Chowdhury *et al*., 2016). Inoculated wild type plants did not show any sign of local callose deposition (**Fig. S5a, b**). Likewise, colonized *vpy-3* mutants showed in general no callose (71%, n=17) (**Fig. S5c**,**d**). However, in 12% of *vpy-3* plants, weak callose accumulation was observed in hypodermal cells with infection hyphae (**Fig. S5e**,**f**), and in 17% of the cases, strong callose accumulation was associated with fungal structures (**Fig. S5g**,**h**). In order to further examine whether callose deposition is related to particular AM fungal fates in the mutants, we performed a second experiment in which all three mutant alleles and the wild type were scored for callose deposition. Wild type plants exhibited a weak level of 4.1% background staining (**Table S1**). Similarly, *vpy* mutants showed weak spurious callose signals in 1.8%-10.1% of the examined infected cells (**Table S1**), and only in 3 cases, a strong callose signal was observed (**Table S1**). These cases did not correlate with particular deformations in the fungus (data not shown) suggesting that there is no correlation between callose formation and fungal abortion.

### Abortion of fungal penetration in petunia *vpy* mutants correlates with local lignin accumulation and induction of lignin biosynthetic genes

We further tested whether the papillae triggered by AM fungal penetration in *vpy* mutants contained lignin. In non-inoculated wild type and mutant roots, a weak general signal in all cell types was observed, except for the stele that showed higher lignin staining (**Fig. 5a**). Penetrated wild type cells occasionally had a weak general signal, but in most cases, no accumulation over background levels was observed (**Fig. 5b**). In contrast, aborted infections in *vpy-1, vpy-2*, and *vpy-3* showed strong local accumulation of lignin in cell wall appositions of the host next to the fungal entry site (**Fig. 5c-e**, arrowheads). Quantification revealed that all three mutant alleles exhibited a significant increase in local lignin accumulation at fungal penetration sites (**Fig. 5f**). For comparison, we evaluated whether the *ram1* mutant also accumulated lignin upon AMF infection (**Fig. S6**). Since *ram1* mutants have a distinct arbuscule phenotype (Rich *et al*., 2015; Rich *et al*., 2017), we first included this developmental stage in the analysis of the wild type. Apart from the lignin signal in the vasculature of the stele (**Fig. S6a**,**b**), infection of the wild type did not lead to appreciable accumulation of lignin around fungal infection coils (**Fig. S6c**,**d**), or in cells with arbuscules (**Fig. S6e**,**f**). Similarly, the infection coils and the defective arbuscules in *ram1* were not accompanied by accumulation of lignin (**Fig. S6g**,**h**; >30 infection and colonization sites assessed, respectively). This indicates that the local accumulation of lignin in *vpy* mutants is a specific phenotypic trait of this mutant and not a general feature of defective symbiosis. Lignin biosynthesis involves a well characterized biosynthetic pathway (Vanholme *et al*., 2019). Thus we identified the respective petunia homologues for all core biosynthetic genes (**Table S3**), and determined their expression during AM development in the wild type and the *vpy* mutants. As expected from lignin stainings (**Fig. 5**), mycorrhizal vpy mutants showed a concerted induction of all lignin biosynthetic genes (**Table 2; Fig. S7; Table S4-S6**).

**Table 2.** Expression ratios of pathogenesis-related (PR) genes in wild type and *vpy* mutants relative to the respective non-mycorrhizal controls. Induction of PR genes was determined by qPCR with actin and glyceraldehyde-3-phosphate dehydrogenase (GAPDH) as reference genes. Values represent -fold induction ratios derived by dividing the expression values of mycorrhizal plants by values of non-mycorrhizal controls (both normalized with the two reference genes) in wt, *vpy-1, vpy-2*, and *vpy-3*. All expression values were derived from 8 biological replicates. Color shading represents induction >2-fold (yellow), >4-fold (orange), and >8-fold (red). Significant induction ratios are indicated by bold type font (two-way ANOVA).

|  | AM(wt)/c(wt) |  | AM(mut)/c(mut) |  |  |
| --- | --- | --- | --- | --- | --- |
|  | AM wt* | AM wt | AM *vpy1* * | AM *vpy2* | AM *vpy3* |
| PR2a | 1.27 | **2.53** | **18.30** | **30.47** | **13.20** |
| PR2b | **2.02** | 1.42 | **8.42** | **17.47** | **24.69** |
| PR2c | 1.48 | 1.26 | 1.87 | 1.09 | 0.88 |
| PR3 | 1.17 | 1.39 | **6.58** | **5.02** | **3.11** |
| PR4b | **2.04** | **5.45** | **9.64** | **41.73** | **13.02** |
| PR4c | 0.92 | 1.10 | **8.71** | **10.76** | **13.82** |
| PR4d | **5.57** | **5.51** | **2.39** | **2.68** | **1.62** |
| PR5a | **6.67** | **3.48** | 1.14 | 1.02 | 1.13 |
| PR6a | 1.80 | **13.24** | 6.11 | **33.76** | 1.57 |
| PR7 | **4.28** | **5.21** | **26.91** | **33.59** | **7.05** |
| PR9 | **6.93** | **5.23** | **24.39** | **6.40** | **26.00** |
| PR14 | **2.53** | **3.02** | **14.91** | **21.59** | **19.79** |
| PR17 | 1.35 | 1.40 | **9.02** | **8.65** | **2.47** |

Yellow: induction>2, orange, induction>4, red: induction>8; bold: p<0.05

**Figure 5.**
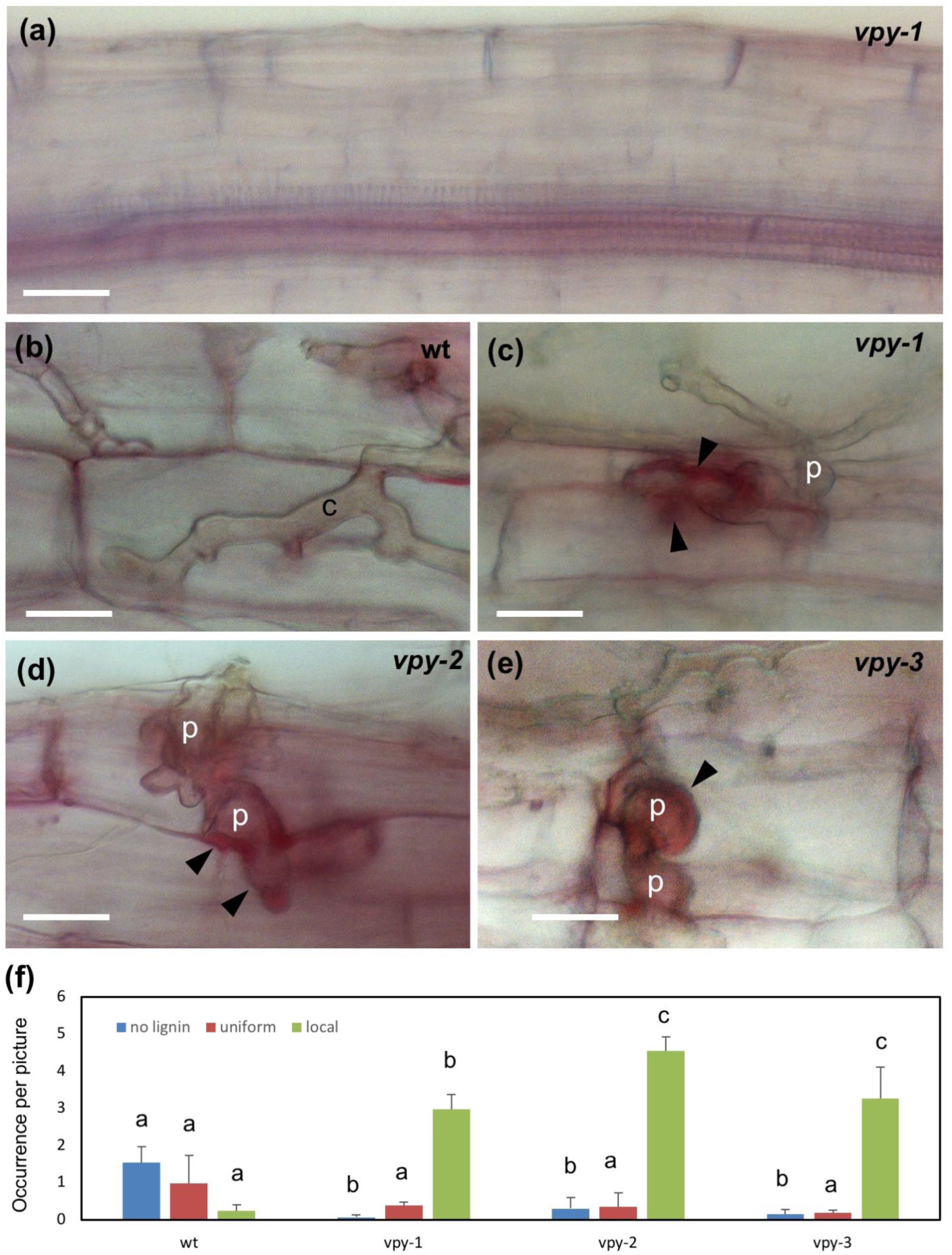
Lignin accumulation in infected hypodermal *vpy* mutant cells. Plants were infected from mycorrhizal nurse plants, and entry points in hypodermal cells of wild type (b), *vpy-1* (c), *vpy-2* (d), and *vpy-3* (e), were evaluated after staining with phloroglucinol-HCl. (a) Non-mycorrhizal *vpy-1* mutant root exhibits weak general signal in all cell types, and elevated signal in the central stele with the vasculature. (b-e) Infected cells of wild type (b), vpy-1 (c), vpy-2 (d), and vpy-3 (e) exhibit weak background signal in the wild type)b), and strong lignin accumulation (arrowheads) in cell wall appositions of *vpy* mutants (c-e). (f) Quantification of lignin accumulation reveals a correlation of strong local lignin accumulation with abortion of AM fungal penetration in hypodermal cells. Size bar, 25 μm.

### Medicago *sym* mutants do not accumulate lignin

In order to assess whether the lignin accumulation phenotype in petunia *vpy* mutants is related to a conserved function of VPY in suppression of defense, we next tested whether *M. truncatula vpy* mutants showed a similar response. Furthermore, we included a number of additional *M. truncatula* mutants with a defect in AM symbiosis (**Table S2**), namely *dmi2-1* (defective in *SYMBIOSIS RECEPTOR-LIKE KINASE, SYMRK*) (Endre *et al*., 2002), *dmi3-1* (defective in *CALCIUM AND CALMODULIN-DEPENDENT PROTEIN KINASE, CCaMK*) (Lévy *et al*., 2004; Mitra *et al*., 2004), *ram1-1* (defective in the *RAM1* orthologue of petunia *RAM1*) (Gobbato *et al*., 2012), and *nsp2-2* (defective in *NODULATION SIGNALLING PATHWAY2, NSP2*) (Kalo *et al*., 2005). As in the case of petunia, the mutants (and the wild type) were inoculated by jive nurse plants to achieve comparable levels of overall root colonization (>50% root length colonization) in all *Medicago* genotypes). Phenotypic assessment of these essentially confirmed all the symbiotic defects described previously (**Fig. S8, S9**). While the wild-type accessions A17 and R108 sustained the formation of hyphal coils in external cells (**Fig. S8a**,**b**), and of arbuscules in the root cortex (**Fig. S8c**), fungal penetration of the root surface was inhibited in *dmi2* and *dmi3* mutants (**Fig. S8d**,**f**). However, due to the strong inoculum potential of nurse plants the fungus eventually entered the roots of both mutant genotypes, however, cortical colonization was markedly different: In *dmi3-1*, only intercellular hyphal colonization was observed (**Fig. S8e**), while in *dmi2-1*, arbuscules were formed at high frequency (**Fig. S8g**), as it was described earlier (Demchenko *et al*., 2004).

*Ram1-1* and *nsp2-2* showed milder phenotypes (**Fig. S9a-d**). In both cases, hyphal coils were formed upon penetration of the root surface (**Fig. S9a**,**c**), however, arbuscule formation was affected in contrasting ways: *ram1-1* mutants only allowed for the formation of small ill-defined arbuscular structures (**Fig. S9b**, arrows)), as described before (Park *et al*., 2015; Rich *et al*., 2015). In contrast, *nsp2-2* contained patchy colonization with apparently normal arbuscules interspersed with arbuscule-free regions or retarded arbuscules (**Fig. S9d**, arrows), implicating nsp2 both, in nodulation as well as in AM (Maillet *et al*., 2011).

Of particular interest was the *Medicago vpy2-2* mutant (**Fig. S9e-h**). *R. irregularis* grew profusely on the root surface and produced complex hyphopodia with many septa (**Fig. S9e**). Occasionally, the root surface was penetrated by thick penetration hyphae (**Fig. S9f**), to reach the cortex where massive intercellular colonization was observed (**Fig. S9g**, arrows). Arbuscules were completely missing, however, intercellular hyphae produced many irregular lateral projections (**Fig. S9h**), presumably representing attempts to invade and colonize cortex cells. Taken together, this pattern of fungal colonization resembles the colonization phenotype of the petunia *vpy* mutant (Sekhara Reddy *et al*., 2007).

We next treated colonized roots of all *Medicago* genotypes with phloroglucinol to reveal lignin deposition. As in petunia, both wild type accessions (A17 and R108) did not accumulate any lignin at the point of fungal infection (**Fig. S10a**,**b**). Surprisingly, however, none of the mutants accumulated significant amounts of lignin at infection points, either (**Fig. S10c-g**), including the *vpy-2* mutant, while prominent lignin deposition was observed along the vasculature in the stele (**Fig. S10h**), as in all other genotypes (data not shown). These results indicate that lignin accumulation at AMF infection points is not part of the phenotype of symbiosis mutants in *Medicago*.

### Resistance of *vpy-3* does not correlate with accumulation of SA, JA, or ethylene

We next tested whether papilla formation in mycorrhizal *vpy* mutants is accompanied by accumulation of the defence hormones salicylic acid (SA), jasmonic acid (JA), or ethylene. In order to assess early infection events, as well as fully established mycorrhizal colonization, we tested *vpy-3* mutant roots after 10 days and 35 days from inoculation. In general, SA levels did not significantly change during AM infection neither in the wild type nor in the mutants (**Fig. S11a**). Next, we compared the levels of free and conjugated SA. SA can be conjugated to sugars or amino acids, resulting in its inactivation. Hence, even though free SA levels may be low after a transient defence response, accumulated SA conjugates can indicate past SA accumulation events. Although conjugated SA occurred in almost 10-fold higher concentrations than free SA, its levels were not induced in the mutant, independent of its mycorrhizal status (**Fig. S11b**). Ethylene was produced only in trace amounts in wild type and *vpy* mutants, and its levels did not change in mutants nor in the wild type upon AM inoculation (data not shown). JA and JA-Ile levels were slightly induced in wild type plants at the second time point, while the levels in mutants were not significantly changed (**Fig. S12**). Again, the JA (**Fig. S12a**) and JA-Ile (**Fig. S12b**) levels were generally low (<1 ppb). Taken together, these results suggest that the cellular resistance reaction in *vpy-3* does not involve the accumulation of SA, ethylene, or JA.

### Induction of PR gene homologues in mycorrhizal roots of wild type and *vpy* mutants

Defence responses often involve induction of defence-related genes, which encode various types of antimicrobial proteins, among them pathogenesis-related (PR) proteins (van Loon *et al*., 2006b). In order to systematically assess the expression of PR genes during mycorrhizal development, we first identified potential PR gene candidates in petunia by protein blasts using established PR protein sequences (mostly from tobacco) as queries (van Loon *et al*., 2006b). For most proteins, at least one predicted close homologue was identified in the *P. axillaris* genome (**Table S7**), and corresponding primer pairs (**Table S7**) were designed for quantitative real time reverse transcription PCR (qRT-PCR) in all three *vpy* alleles.

Consistent with previous findings (Salzer *et al*., 2000; Breuillin *et al*., 2010; Campos-Soriano *et al*., 2010), several PR gene homologues were induced in mycorrhizal roots of wild type plants (**Table 2**), in particular ß-1,3-glucanase (PR2a), chitinase (PR4b and PR4d), a thaumatin-like gene (PR5a), a proteinase inhibitor (PR6a), a proteinase (PR7), a peroxidase (PR9), and a lipid transfer protein (PR14). In *vpy* mutants, expression of several genes was further induced (**Table 2**), and some were induced only in the mutants, namely PR2b (ß-1,3-glucanase), PR3 (chitinase), PR4c (chitinases), and PR17 (unknown function). Comparing directly mycorrhizal mutants with mycorrhizal wild type plants showed which genes are particularly induced in the mutants (PR2a, PR2b, PR4c; **Table S8** left), and comparison of non-mycorrhiozal mutants and wild type showed that the mutants had constitutively higher levels of PR4d and PR6a expression (**Table S8** right).

In order to identify PR genes that are responsive to MAMPs, we treated non-mycorrhizal wild type plants with a chitin hydrolysate that contains N-acetyl-glucosamine oligosaccharides (Chit), and an aqueous extract from *Penicillium chrysogenum* (Pen). Both preparations had previously been shown to induce a strong defence response and disease resistance in *Arabidopsis thaliana* (Thuerig *et al*., 2005). Several PR genes were induced by Pen elicitor, only two of them were responsive to chitin (**Table S9**). The strongest induction was found for PR4d, one of the two genes that is constitutively induced in *vpy* mutants (**Table S8**).

Taken together, these results indicate that i.) wild type mycorrhizal roots express several PR genes at induced levels, ii.) mycorrhizal *vpy* mutants are overresponsive with regard to AM-inducible PR genes, and they induce additional PR genes, and iii.) several PR genes are responsive to MAMPS in elicitor preparations.

### Cell wall appositions in *vpy-3* contain ß-1,3-glucanase

PR proteins may accumulate locally at sites of microbial infection, or they could be generally induced without a particular pattern of accumulation. Since ß-1,3-glucanase was induced in *vpy* mutants (PR2a and PR2b in **Table 2**), we investigated whether papillae in *vpy* mutants contain ß-1,3-glucanase by employing a polyclonal antiserum raised against tobacco ß-1,3-glucanase (Beffa *et al*., 1993). Immuno-cytochemical detection with gold-coupled secondary antibodies revealed a low generally distributed signal in the cell walls of non-inoculated control roots in the wild type as in *vpy-3* mutants (**Fig. S13a, c, e, g**). Inoculated wild type plants did not show an obvious increase in the vicinity of fungal hyphae (**Fig. S13b**,**f**), consistent with the weak induction of PR2 in mycorrhizal wild type plants (**Table 2**). Inoculated *vpy-3* mutants exhibited a general distribution of gold particles at similar levels as the other treatments (**Fig. S13d**,**h**). However, papillae in the vicinity of aborted fungal hyphae exhibited a more prominent signal (**Fig. S14, S15**). Hence, accumulation of ß-1,3-glucanase appears to increase in papillae of *vpy-3*. Quantification of immunogold signal confirmed the local accumulation of ß-1,3-glucanase in papillae (Fig. S16).

## DISCUSSION

### Signaling in symbiosis and defence

Transcript profiling of AM in various host species has shown that besides the induction of many AM-related genes with presumed functions in symbiosis (e.g. phosphate and ammonium transporters), many AM-induced genes are related to established defense genes (e.g. PR genes) (Gao *et al*., 2004; Grunwald *et al*., 2004; Deguchi *et al*., 2007; Liu *et al*., 2007; Siciliano *et al*., 2007; Fiorilli *et al*., 2009; Breuillin *et al*., 2010; Campos-Soriano *et al*., 2010; Gaude *et al*., 2012; Handa *et al*., 2015; Fiorilli *et al*., 2018). In addition, we know that the complex signaling mechanism in symbiosis includes, besides dedicated symbiosis signals (myc factors) molecular species that are known to also play a role in disease resistance, e.g. chitin oligomers (Zipfel & Oldroyd, 2017). Consistent with these findings, some receptors for chitinous signaling molecules have a dual role in disease resistance and in symbiosis (Zipfel & Oldroyd, 2017). These findings point to overlapping mechanisms in the signaling pathways in symbiosis and defense, including for example the production of ROS (Scheler *et al*., 2013; Damiani *et al*., 2016), the induction of JA (Wasternack & Hause, 2013), and calcium-related signals (Aldon *et al*., 2018), which all have been observed in both, symbiosis and defense. Moreover, the outcome of AM symbiosis for the host plant is highly context-dependent (Klironomos, 2003; Smith *et al*., 2009; Lanfranco *et al*., 2018). This implies that some additional mechanisms may be required to specify and determine whether roots engage either in defense or in symbiosis. It is conceivable that the CSSP is one of the central elements to ensure that symbiosis-related signaling overrides defense signalling during mycorrhizal infection. In addition to specific symbiosis signals, AMF use effectors to help prevent the induction of defense responses, or to dampen their amplitude (Kloppholz *et al*., 2011; Sedzielewska Toro & Brachmann, 2016; Tang *et al*., 2016; Kamel *et al*., 2017; Chen, ECH *et al*., 2018; Voss *et al*., 2018; Morin *et al*., 2019).

### Symbiosis mutants show symptoms of a defense response

Consistent with the assumption that commitment to symbiosis requires repression of defense reactions, many mutants that are defective in symbiosis signaling (*sym* mutants) show various symptoms of a cellular defense response (Gollotte *et al*., 1993; Gianinazzi-Pearson, 1996; Wegel *et al*., 1998; Ruiz-Lozano *et al*., 1999; Bonfante *et al*., 2000; Marsh & Schultze, 2001; Novero *et al*., 2002; Demchenko *et al*., 2004). In such mutants, AMF colonization is hampered or fully blocked, cells accumulate secondary metabolites that are known (or suspected) to act as defense agents, cell walls are altered or reinforced, and defense-related transcripts are induced. This is strong evidence that one function of symbiotic signaling is to avoid, or repress, defense during AM interactions.

In *vpy* mutants, defense-like aspects of the AM-defective phenotype are particularly prominent. Here, we show that this syndrome resembles a *bona-fide* cellular defense response, involving the formation of cell wall papillae (**Figs. 2-3; Fig. S1-S2**), the local accumulation of lignin (**Fig. 5**), the induction of lignin-biosynthetic genes (**Table 1**), and the induction of a wide array of PR genes (**Table 2**). The fact that *vpy* mutants in petunia show these symptoms of defense, suggests that one function of VPY is to suppress, directly or indirectly, the induction of defense. This appears to be a prerequisite for intracellular accommodation of AMF during infection at the root surface, and during arbuscule development in the cortex (Ercolin & Reinhardt, 2011; Gutjahr & Parniske, 2013). An interesting aspect of the colonization phenotype of *vpy* mutants in both, petunia and *Medicago*, is that intercellular hyphal colonization was not inhibited (**Fig. 2a, Fig. S9g**) (Sekhara Reddy *et al*., 2007), suggesting that the interaction at the extracellular level is compatible. A similar phenotype was observed in *della* mutants that were essentially devoid of intracellular structures, yet exhibited high colonization levels by profusely growing intercellular hyphae (Floss *et al*., 2013).

### Defense response in petunia *vpy* mutants involves local lignin accumulation

Papilla formation is considered a highly effective defence mechanisms of plants against fungal pathogens (Hückelhoven, 2007). Papillae inhibit fungal penetration of aerial plant tissues and often result in complete abortion of the pathogen. In the root, the formation of cell wall reinforcements as defence mechanism has been less explored. The cell wall appositions in mycorrhizal *vpy* mutants colonized by *R. irregularis* were either locally restricted to the site of cell penetration, or they surrounded intracellular hyphae thereby encapsulating them (**Fig. 2, 3, 5, S1, S2, S14**).

Interestingly, the papillae of *vpy* mutants did in most cases not contain callose, and the cell walls were not autofluorescent, indicating that some hallmarks of defence are absent in mycorrhizal *vpy* mutants. However, papillae were impregnated by local accumulation of lignin (**Fig. 5**). Consistent with this observation, the lignin biosynthetic pathway was induced in a concerted fashion, in inoculated *vpy* mutants, involving all biosynthetic genes (**Fig. S7**) (Vanholme *et al*., 2019). Localized cell wall lignification is among the most effective cellular defence responses of plants (Miedes *et al*., 2014). Hence the correlation of cell wall reinforcement and localized lignin impregnation indicates that lignin may contribute to the abortion of AM fungal cell penetration in petunia *vpy* mutants. In addition, the concerted induction of enzymes in phenylpropanoid pathway (**Table 1**; **Fig. S7**) could potentially lead to the production of other defense related compounds (Dixon *et al*., 2002; Vogt, 2010). Interestingly, the *ram1* mutant which exhibits a later defect in AM development during arbuscule formation, and which does not display markers of a cellular defense response (Park *et al*., 2015; Rich *et al*., 2015; Pimprikar *et al*., 2016) did not accumulate any lignin (**Fig. S6**). Strikingly, in *M. truncatula vpy* mutants (as well as *dmi2-1, dmi3-1, nsp2-2*, and *ram1-1* mutants), no accumulation of lignin was detected, suggesting that in *M. truncatula* roots, abortion of AMF infection does not involve lignin. This points to taxon-specific aspects in root defense, and perhaps in AM-related pathways between petunia and *M. truncatula*, and perhaps more generally, between Solanaceae and legumes.

### Induction of PR genes in mycorrhizal wild type and *vpy* mutants

A characteristic symptom of defence in plants is the accumulation of pathogenesis-related (PR) proteins (van Loon *et al*., 2006b). Several PR proteins have been shown to have antimicrobial activity, and some of them contribute to disease resistance, hence they are regarded as markers of defence. We assessed the expression of all PR gene homologues that could be identified in the genome of *P. axillaris*, the parent that contributed ca. 80% to the *P. hybrida* genome (Bombarely *et al*., 2016). From a list of 17 PR genes (van Loon *et al*., 2006b), 13 were found to be expressed in roots (**Table 2**). PR genes have previously been reported to be induced in many mycorrhizal associations (reviewed in (García-Garrido & Ocampo, 2002). In most cases, the induction was characterized as early, weak and transient (Gianinazzi-Pearson *et al*., 1996; Kapulnik *et al*., 1996). However, in some cases, PR genes were induced in a sustained fashion and at high levels (Salzer *et al*., 2000; Breuillin *et al*., 2010), indicating that they may have an important role in AM symbiosis. In addition, the phylogenomic signature of some AM-induced PR proteins, e.g. chitinase III, indicates that they have been under positive selection in AM-competent plant species (Rich *et al*., 2014), suggesting that they have specific functions in AM.

We observed that several PR genes were induced in mycorrhizal wild type roots (**Table 2**), possibly reflecting an elevated defence status in mycorrhizal roots (García-Garrido & Ocampo, 2002; Pozo & Azcon-Aguilar, 2007). In mycorrhizal *vpy* mutant roots, several PR genes (PR2a, PR2b, PR4c, PR7, PR9, PR14, and PR17) were expressed at even much higher levels than in the wild type, consistent with our findings that mycorrhizal *vpy* mutants mount a defense response (**Fig. 2-5**). The strong induction of two PR2 genes in *vpy* mutants, and the local accumulation in papillae of PR2 protein (**Fig. S10**), are of particular interest since transgenic tobacco overexpressing PR2 exhibited significantly delayed fungal colonization of the roots (Vierheilig *et al*., 1995).

### Defence response in *vpy* mutants does not affect levels of SA, JA, ethylene or ROS

Initiation of a defence response in plants is often associated with the accumulation of the stress and defence hormones salicylic acid (SA), jasmonic acid (JA), or ethylene (Browse, 2009; Vlot *et al*., 2009; Broekgaarden *et al*., 2015). In mycorrhizal roots, the picture is more complex, since SA and JA can be either up- or downregulated during symbiosis (Fernandez *et al*., 2014). We found no significant differences in the levels of free or conjugated SA (**Fig. S11**), consistent with the lack of induction of the SA-marker PR1. JA levels were not affected at the early time point of the interaction (10d), but later on, the wild type exhibited slightly higher levels than *vpy-3* (**Fig. S12**). JA levels are known to be induced in mycorrhizal roots in some cases, however, the role of JA in symbiosis is controversial (Wasternack & Hause, 2013).

A commonly observed phenomenon associated with plant defense is the production of reactive oxygen species (ROS) in the host, also known as the oxidative burst (Torres *et al*., 2006). In AM, ROS may also play a role, although in this case, it is not clear which of the symbiotic partners is the main source of ROS (Fester & Hause, 2005). We used two established staining procedures to detect H_2_O_2_ (DAB) and O_2_^-^ (NBT), respectively. Interestingly, both methods revealed high ROS levels in the fungus, irrespective of the host genotype (**Fig. 4, S3, S4**). As described in *M. truncatula* (Salzer *et al*., 1999), highest H_2_O_2_ levels were associated with clumped arbuscules that appeared to undergo senescence (**Fig. S3**). The resolution of the technique does not allow to assign the DAB signal with certainty to either the fungus or the host in cells with arbuscules, while in the case of fungal vesicles, the DAB signal was clearly confined to the fungal cytoplasm. NBT staining in the host was strongest in root tips (**Fig. S4a-e**), as it has been shown for *Arabidopsis* and *Medicago* (Dunand *et al*., 2007; Chen *et al*., 2015). In infected areas of the cortex, NBT signal was confined to fungal hyphae (**Fig. S4f-k**) and arbuscules (in the wild type). Taken together, these results show that none of the classical defense hormones (SA, JA, ethylene) were induced during the defense response in *vpy-3*, and ROS accumulation patterns did not reveal a correlation with defence in *vpy* mutants.

### VAPYRIN is involved in repression of cellular defence

How could VPY interfere with defence? The localization of VPY to mobile subcellular compartments (Feddermann *et al*., 2010; Pumplin *et al*., 2010), and its interaction with EXO70I at the tip of hyphal branches (Zhang *et al*., 2015) suggest that VPY is involved in subcellular trafficking towards fungal hyphae. Hence, the mobile compartments could transport a cargo or a membrane component that interferes with the induction of defence. Alternatively, VPY could be a target of an AM fungal effector which represses defence in the host. Future research should identify the interaction partners of VPY and potential downstream components to explain how it acts on defence mechanisms.

## Acknowledgements

We thank Christine Lang and Eliane Abou-Mansour for assistance with salicylic acid determination, and Jürg Felix for providing chitin and Penicillium MAMP preparations.

## Author contributions

MC, SB, LB, GD, MS, and DR planned and conducted the experiments and were involved in data collection. GG performed hormone analytics. SB, MC, and DR were involved in data analysis. DR wrote the paper with assistance of SB and MC.

## Supplementary material

**Figure S1. Confocal analysis of papilla formation in *vpy* mutants**.

Wild type (a), *vpy-1* (b), *vpy-2* (c), and *vpy-3* (d), were harvested 33 days after nurse plant inoculation, stained with basic fuchsin and wheat germ agglutinine (WGA)-Alexa488, and evaluated for cell wall appositions by confocal microscopy. Shown are hyphae (green) and cell walls with papillae (red). Papillae are indicated by arrowheads. (e) Quantification of papilla formation. Representative pictures of penetrated cells as in (a)-(d) were assessed for the number of papillae. Shown are the mean + Stdv of three samples containing at least 8 pictures each. Significance of differences was tested by two-way ANOVA. Size bar 25 μm.

**Figure S2. Infection phenotype in *vpy-3* root cortex cells**.

Transmission electron micrographs of wild type (a-c,f), and *vpy-3* (d,e) root cortex cells after infection from mycorrhizal nurse plants.

(a) An intercellular hypha (IH) has produced an intracellular penetration hypha (PH; arrowheads), and several thin-walled fine fungal arbuscule branches that are embedded in the cytoplasm of the host (asterisks). The space between the fungal cell wall and the periarbuscular membrane of the host is very thin.

(b) Fully colonized cortex cell containing an arbuscule. Note large trunk hyphae (TH) and thin arbuscular branches (asterisks).

(c) Cortex cell with a senescing arbuscule exhibiting collapsed fungal material (arrows). As in (b), the space between the fungal cell walls and the periarbuscular membrane is very narrow, and no plant derived cell wall appositions can be seen around fungal material.

(d) Attempted entry into a *vpy-3* cortex cell results in formation of a papilla (arrowheads).

(e) Branched fungal structure in the cortex of a *vpy-3* root. In the vicinity of the fungal hyphae (green), multiple small papillae (red, arrowheads) were formed.

(f) Two arbuscules in the cortex of a wild type plant. Fungal hyphae (green) did not elicit any papillae.

(a)-(d) electron micrographs; (e,f) Confocal micrographs.

Scale bars in (a-d), 2 μm; in (e,f) 25 μm.

PH, penetration hypha; TH, trunk hypha; x, xylem; IH, intercellular hypha

**Figure S3. Cytochemical detection of H**_**2**_**O**_**2**_ **in the cortex of *vpy* mutants**

Roots of all *vpy* alleles were harvested four weeks after nurse plant inoculation and evaluated after staining with 3,3’-diaminobenzidine (DAB) to detect H_2_O_2_. Shown are representative DAB-stained *vpy-2* mutant roots (a-c), and corresponding wild type roots (d-f). All three *vpy* alleles gave similar results.

(a) Weak background staining in *vpy-2*. The other *vpy* mutants and the wild type showed a similar background signal.

(b) Intermediate signal associated with residual intracellular colonization in cortex cells (asterisks).

(c) Strong signal in vesicles (v).

(d) In wild type roots, strong staining was observed in arbuscules (asterisks).

(e) Staining was further increased in cells with clumping arbuscules (double asterisks), relative to mature arbuscules (asterisk).

(f) Strong signal in vesicles (v) in the wild type cortex. Size bar 50 µm.

**Figure S4. Cytochemical detection of O**_**2**_^**-**^ **in *vpy* mutants**

Wild type and *vpy* mutants were harvested four weeks after nurse plant inoculation and evaluated after staining with nitroblue tetrazolium (NBT) to detect O_2_^-^. Strongest staining was observed in root tips (a-d) and lateral roots (e) of all genotypes (as indicated), independent of AM inoculation. In colonized areas of all genotypes (as indicated), strongest signal was detected in all fungal structures (f-j). White arrowheads indicate intercellular longitudinal hyphae; yellow arrowheads mark their tips, black arrowheads point to cell wall thickenings in hypodermal cells. m, root meristem; p, penetration hypha. Size bars 100 µm in (a-e), 25 µm in (f-j).

**Figure S5. Callose accumulation associated with aborted fungal infection of *vpy-3***. Plants were infected from mycorrhizal nurse plants, and entry points in hypodermal cells of wild type (a,b), and *vpy-3* (c-h), were evaluated after staining with aniline blue. (a,c,e,g) Bright field images. (b,d,f,h) Epifluorescence images of the same cells.

(a,b) Successful penetration of a wild type hypodermal cell. No callose can be observed around the infection hypha (asterisk).

(c,d) Penetrated cell of a *vpy-3* mutant root with a deformed infection hypha (asterisk) but no callose deposition.

(e,f) Aborted fungal entry point into a hypodermal cell (asterisk) with weak callose accumulation (arrowheads).

(g,h) Aborted fungal entry into a hypodermal cell (asterisk) with the accumulation of callose-rich cell wall material (arrowhead). Scale bars, 20 μm.

**Figure S6. Lack of lignin accumulation in mycorrhizal *ram1* mutants**

Non-colonized roots (a,b) or roots of colonized wild type (c-f), or colonized *ram1* mutants (g,h) were processed and stained with Trypan Blue (b, d, f, h) or phloroglucinol (a, c, e, g), respectively. (a, b) non-colonized cells, (c,d) intracellular hypodermal coils, (e,f) arbuscules in the wild type, (g,h) infection unit with a hyphal coil (c), and aberrant hyphal colonization in the cortex (arrows). vasc, vascular bundles in the stele; HPC, hypodermal passage cell; c, hyphal coil in hypodermal cells; a, arbuscules. size bars 25 µm.

**Figure S7. Figure 1. Concerted induction of lignin biosynthetic genes in mycorrhizal *vpy* mutants**. <u>The main biosynthetic route toward the monolignols p-coumaryl, coniferyl, and sinapyl alcohol is shown. Red numbers indicate the induction ratios of the respective gene functions</u> calculated by averaging the induction ratios for the individual lignin-biosynthetic genes in a functional family, and by averaging the three *vpy* mutants. Pathway scheme after {Vanholme, 2010 #6704}; <u>see Table 1).</u>

<u>PAL, PHENYLALANINE AMMONIA-LYASE;</u>

<u>C4H, CINNAMATE 4-HYDROXYLASE;</u>

<u>4CL, 4-COUMARATE:CoA LIGASE;</u>

<u>C3H, p-COUMARATE 3-HYDROXYLASE;</u>

<u>HCT, p-HYDROXYCINNAMOYL-CoA:QUINATE/ SHIKIMATE p-</u>

<u>HYDROXYCINNAMOYLTRANSFERASE;</u>

<u>CCoAOMT, CAFFEOYL-CoA O-METHYLTRANSFERASE;</u>

<u>CCR, CINNAMOYL-CoA REDUCTASE;</u>

<u>F5H, FERULATE 5-HYDROXYLASE;</u>

<u>COMT, CAFFEIC ACID O-METHYLTRANSFERASE;</u>

<u>CAD, CINNAMYL ALCOHOL DEHYDROGENASE.</u>

**Figure S8. Mycorrhizal colonization pattern in *M. truncatula sym* mutants** Colonized roots were stained with Trypan Blue. (a-c) wild type accessions A17 and R108 (as indicated). (d,e) *dmi3-1* mutant (defective in *CCaMK*), (f,g) *dmi2-1* mutant (defective in *SYMRK*). (a,b) Hyphal coils in hypodermal cells, (c) arbuscules in the cortex, (d,f) fungal hyphae growing on the root surface with attempted entry points with hyphopodia (asterisks), (e,g) cortical colonization pattern with intercellular hyphae (arrows) (e), and intracellular arbuscules (g). a, arbuscules; c, hyphal coils. Size bars 25 µm.

**Figure S9. Mycorrhizal colonization pattern in *M. truncatula sym* mutants** Colonized roots were stained with Trypan Blue. (a,b) *ram1* mutants; (c,d) *nsp2-2* mutants; (e-h), *vpy-2* mutants. (a,c) Entry point with hyphal coils; (b) Aberrant arbuscules (arrows) in the *ram1* mutant; (d) widely spaced normal arbuscules with intervening partially developed structures (arrows). (e) Hyphae growing on the root surface with hyphal swellings resembling hyphopodia (asterisks); (f) Fungal entry point with a thick penetration hypha (p); (g,h) profusely growing intercellular hyphae (arrows), occasionally with lateral projections that may represent attempted cell invasion events. a, arbuscules; c, hyphal coils; p, penetration peg. Size bars 25 µm.

**Figure S10. Lack of lignin accumulation at fungal entry points in *M. truncatula sym* mutants**

Colonized roots were stained with phloroglucinol. (a,b) Wild type accessions A17 and R108, respectively. (c) *dmi3-1* mutant with attempted infection points represented by hyphal swellings (asterisks), and accumulation of brownish material in hosts cells (arrows); (d) *dmi2-1* with attempted infection points represented by hyphal swellings (asterisks), and accumulation of brownish material next to the fungal hyphae (arrow); (e) Hyphal coil in a hypodermal cell of a *ram1-1* mutant; (f) Hyphal coil in a hypodermal cell of an *nsp2-2* mutant; (g) Entry point in a vpy-2 mutant by a hyphal penetration peg (p), and subsequent intercellular hypha; (h) Non-colonized root showing staining of the vasculature in the stele. c, hyphal coils; p, penetration peg; vasc, vasculature. Size bars 25 µm.

**Figure S11. Salicylic acid levels in mycorrhizal wild type and *vpy-3* roots**.

(a) Accumulation of free salicylic acid in wild type *P. hybrida* (white columns), and *vpy-3* plants (black columns) after 10 days (10d) and 35 days (35d) with or without inoculation with *R. irregularis* (AM).

(b) Free SA (black columns) and conjugated SA (white columns) in wild type and vpy-3 roots with or without inoculation with *R. irregularis* (AM).

Columns represent the average of 5 biological replicates +Stdv. There was no significant difference between any of the treatments (student’s T-test).

**Figure S12. Jasmonic acid levels in mycorrhizal wild type and *vpy-3* roots**.

(a) Accumulation of free jasmonic acid in wild type *P. hybrida* (white columns), and *vpy-3* plants (black columns) after 10 days (10d) and 35 days (35d) with or without inoculation with *R. irregularis* (AM).

(b) Accumulation of isoleucine-conjugated jasmonic acid (JA-Ile) in wild type *P. hybrida* (white columns), and *vpy-3* plants (black columns) after 10 days (10d) and 35 days (35d) with or without inoculation with *R. irregularis* (AM). Columns represent the average of 5 biological replicates +Stdv. Wild type plants, but not *vpy-3* mutants, showed a significant increase of JA-Ile in mycorrhizal roots, relative to non-mycorrhizal controls (student’s T-test).

**Figure S13. Immunocytochemical analysis of β-1**,**3-glucanase in mycorrhizal roots**.

Hypodermal cells (a-d), and cortical cells (e-h) of wild type (a,b,e,f), and *vpy-3* mutant roots (c,d,g,h). Non-inoculated control plants (a,c,e,g) and mycorrhizal plants (b,d,f,h) were analyzed by immunogold staining with antibodies raised against tobacco β-1,3-glucanase. Low levels of immunogold were detected in hypodermal cells (a-d), while label was almost undetectable in cortex cells (e-h). va, plant vacuole; hl, hypodermal layer; 1°, primary cell wall; 2°, secondary cell wall; fw, fungal cell wall. Scale bars, 250 nm.

**Figure S14. Accumulation of β-1**,**3-glucanase in a cell wall apposition of *vpy-3***. Same sample as in Fig. 8d. An aborted fungal penetration hypha (degenerating fungus) grown next to a hypodermal cell (upper right) and a cortex cell (lower left). The hypodermal cell has formed a thick cell wall apposition that exhibits strong immunogold signal. hl, hypodermal layer; 1°, primary cell wall; 2°, secondary cell wall. Scale bar, 500 nm.

**Figure S15. Low-magnification overview picture of the sample shown in Fig. S9**.

**Figure S16. Quantification of immunogold signal of representative pictures as shown in Fig. 7 and Fig. 8**.

**Table S1. Quantification of callose accumulation in representative samples as shown in Fig. S5**.

**Table S2. *M. truncatula* mutants used in this study**

**Table S3. Primers used for qPCR of lignin-related genes**

**Table S4. Induction of lignin-related genes in *vpy* mutants vs. wild type either in the mycorrhizal or non-mycorrhizal context**.

Induction of lignin biosyntheteic genes was determined by qPCR with actin and glyceraldehyde-3-phosphate dehydrogenase (GAPDH) as reference genes. Values represent -fold induction ratios derived by dividing the expression values of mutants by the values from the wild type, either in mycorrhizal (left) or control conditions (right). All expression values represent the average of 8 biological replicates. Color shading represents induction >2-fold (yellow), >4-fold (orange), and >8-fold (red). Significant induction ratios are indicated by bold type font (two-way ANOVA).

**Table S5. Expression values for lignin-related genes for the experiment on *vpy-2* and *vpy-3* from the genes for which induction ratios were calculated in Table 1 and Table S3**.

**Table S6. Expression values for lignin-related genes for the experiment on *vpy-1* from the genes for which induction ratios were calculated in Table 1 and Table S3**.

**Table S7. Primers used for qPCR of PR genes**

**Table S8. Induction of pathogenesis-related (PR) genes in mutants vs. wild type either in the mycorrhizal or non-mycorrhizal context**.

Induction of PR genes was determined by qPCR with actin and glyceraldehyde-3-phosphate dehydrogenase (GAPDH) as reference genes. Values represent -fold induction ratios derived by dividing the expression values of mutants by the values from the wild type, either in mycorrhizal (left) or control conditions (right). All expression values represent the average of 8 biological replicates. Color shading represents induction >2-fold (yellow), >4-fold (orange), and >8-fold (red). Significant induction ratios are indicated by bold type font (two-way ANOVA).

**Table S9. Induction of pathogenesis-related (PR) genes in wild type plants treated with fungal elicitor preparations**.

Induction of PR genes was determined by qPCR with actin and glyceraldehyde-3-phosphate dehydrogenase (GAPDH) as reference genes. Values represent normalized expression values from control roots, or from roots treated with chitin hydrolysate (Chit), or Penicillium preparation (Pen) after 1h or 4h of treatment. All expression values represent the average of 8 biological replicates. Color shading represents induction >2-fold (yellow), and >4-fold (orange). Significant induction ratios are indicated by bold type font (two-way ANOVA).

**Table S10. Expression values for the experiment on *vpy-2* and *vpy-3* from the genes for which induction ratios were calculated in Table 1 and Table S3**.

**Table S11. Expression values for the experiment on *vpy-1* from the genes for which induction ratios were calculated in Table 1 and Table S3**.

**Table S12. Expression values from the experiment wild type plants treated with elicitors from genes for which induction ratios were calculated in Table S4**.

## References

Aldon D, Mbengue M, Mazars C, Galaud JP. 2018. Calcium signalling in plant biotic interactions. International Journal of Molecular Sciences 19(3).

Altman LG, Schneider BG, Papermaster DS. 1984. Rapid embedding of tissues in Lowicryl K4M for immunoelectron microscopy. Journal of Histochemistry and Cytochemistry 32(11): 1217–1223.

Bapaume L, Laukamm S, Darbon G, Monney C, Meyenhofer F, Feddermann N, Chen M, Reinhardt D. 2019. VAPYRIN marks an endosomal trafficking compartment involved in arbuscular mycorrhizal symbiosis. Frontiers in Plant Science 10.

Beffa RS, Neuhaus JM, Meins F. 1993. Physiological compensation in antisense transformants: Specific induction of an “ersatz”-glucan endo-1,3-ß-glucosidase in plants infected with necrotizing viruses. Proceedings Of The National Academy Of Sciences Of The United States Of America 90(19): 8792–8796.

Boller T, Felix G. 2009. A renaissance of elicitors: Perception of microbe-associated molecular patterns and danger signals by pattern-recognition receptors. Annual Review Of Plant Biology 60: 379–406.

Bombarely A, Moser M, Consortium TP. 2016. Insight into the evolution of the Solanaceae from the parental genomes of *Petunia hybrida*. Nature Plants 2: 16074–16082.

Bonfante P, Genre A, Faccio A, Martini I, Schauser L, Stougaard J, Webb J, Parniske M. 2000. The *Lotus japonicus LjSym4* gene is required for the successful symbiotic infection of root epidermal cells. Molecular Plant-Microbe Interactions 13(10): 1109–1120.

Breuillin F, Schramm J, Hajirezaei M, Ahkami A, Favre P, Druege U, Hause B, M. B, Kretzschmar T, Bossolini E, et al. 2010. Phosphate systemically inhibits development of arbuscular mycorrhiza in *Petunia hybrida* and represses genes involved in mycorrhizal functioning. Plant Journal 64: 1002–1017.

Broekgaarden C, Caarls L, Vos IA, Pieterse CMJ, Van Wees SCM. 2015. Ethylene: Traffic controller on hormonal crossroads to defense. Plant Physiology 169(4): 2371–2379.

Browse J. 2009. Jasmonate passes muster: A receptor and targets for the defense hormone. Annual Review Of Plant Biology 60: 183–205.

Campos-Soriano L, Garcia-Garrido M, San Segundo B. 2010. Activation of basal defense mechanisms of rice plants by *Glomus intraradices* does not affect the arbuscular mycorrhizal symbiosis. New Phytol 188: 597–614.

Chen DS, Liu CW, Roy S, Cousins D, Stacey N, Murray JD. 2015. Identification of a core set of rhizobial infection genes using data from single cell-types. Frontiers in Plant Science 6.

Chen ECH, Morin E, Beaudet D, Noel J, Yildirir G, Ndikumana S, Charron P, St-Onge C, Giorgi J, Krüger M, et al. 2018. High intraspecific genome diversity in the model arbuscular mycorrhizal symbiont *Rhizophagus irregularis*. New Phytologist 220(4): 1161–1171.

Chen M, Arato M, Borghi L, Nouri E, Reinhardt D. 2018. Beneficial services of arbuscular mycorrhizal fungi - From ecology to application. Frontiers in Plant Science 9.

Chowdhury J, Henderson M, Schweizer P, Burton RA, Fincher GB, Little A. 2014. Differential accumulation of callose, arabinoxylan and cellulose in nonpenetrated versus penetrated papillae on leaves of barley infected with *Blumeria graminis* f. sp *hordei*. New Phytologist 204(3): 650–660.

Chowdhury J, Schober MS, Shirley NJ, Singh RR, Jacobs AK, Douchkov D, Schweizer P, Fincher GB, Burton RA, Little A. 2016. Down-regulation of the *glucan synthase-like 6* gene (*HvGsl6*) in barley leads to decreased callose accumulation and increased cell wall penetration by *Blumeria graminis* f. sp hordei. New Phytologist 212(2): 434–443.

Chuberre C, Plancot B, Driouich A, Moore JP, Bardor M, Gugi B, Vicre M. 2018. Plant immunity is compartmentalized and specialized in roots. Frontiers in Plant Science 9.

Damiani I, Pauly N, Puppo A, Brouquisse R, Boscari A. 2016. Reactive Oxygen Species and Nitric Oxide Control Early Steps of the Legume - Rhizobium Symbiotic Interaction. Frontiers in Plant Science 7.

Daudi A, O’Brien JA. 2016. Detection of hydrogen peroxide by DAB staining in arabidopsis leaves. Bio Protocols 2012(2): 18.

David R, Itzhaki H, Ginzberg I, Gafni Y, Galili G, Kapulnik Y. 1998. Suppression of tobacco basic chitinase gene expression in response to colonization by the arbuscular mycorrhizal fungus *Glomus intraradices*. Molecular Plant-Microbe Interactions 11(6): 489–497.

Deguchi Y, Banba M, Shimoda Y, Chechetka SA, Suzuri R, Okusako Y, Ooki Y, Toyokura K, Suzuki A, Uchiumi T, et al. 2007. Transcriptome profiling of Lotus japonicus roots during arbuscular mycorrhiza development and comparison with that of nodulation. Dna Research 14(3): 117–133.

Demchenko K, Winzer T, Stougaard J, Parniske M, Pawlowski K. 2004. Distinct roles of *Lotus japonicus SYMRK* and *SYM15* in root colonization and arbuscule formation. New Phytologist 163(2): 381–392.

Dixon RA, Achnine L, Kota P, Liu CJ, Reddy MSS, Wang LJ. 2002. The phenylpropanoid pathway and plant defence - a genomics perspective. Molecular Plant Pathology 3(5): 371–390.

Dunand C, Crevecoeur M, Penel C. 2007. Distribution of superoxide and hydrogen peroxide in *Arabidopsis* root and their influence on root development: possible interaction with peroxidases. New Phytologist 174(2): 332–341.

Endre G, Kereszt A, Kevei Z, Mihacea S, Kaló P, Kiss GB. 2002. A receptor kinase gene regulating symbiotic nodule development. Nature 417(6892): 962–966.

Ercolin F, Reinhardt D. 2011. Successful joint ventures of plants: arbuscular mycorrhiza and beyond. Trends In Plant Science 16(7): 356–362.

Feddermann N, Duvvuru Muni RR, Zeier T, Stuurman J, Ercolin F, Schorderet M, Reinhardt D. 2010. The *PAM1* gene of petunia, required for intracellular accommodation and morphogenesis of arbuscular mycorrhizal fungi, encodes a homologue of VAPYRIN. Plant Journal 64(3): 470–481.

Feddermann N, Reinhardt D. 2011. Conserved residues in the ankyrin domain of VAPYRIN indicate potential protein-protein interaction surfaces. Plant Signaling and Behavior 6(5): 680–684.

Fernandez I, Merlos M, Lopez-Raez JA, Martinez-Medina A, Ferrol N, Azcon C, Bonfante P, Flors V, Pozo MJ. 2014. Defense related phytohormones regulation in arbuscular mycorrhizal symbioses depends on the partner genotypes. Journal Of Chemical Ecology 40(7): 791–803.

Fester T, Hause G. 2005. Accumulation of reactive oxygen species in arbuscular mycorrhizal roots. Mycorrhiza 15(5): 373–379.

Fiorilli V, Catoni M, Miozzi L, Novero M, Accotto GP, Lanfranco L. 2009. Global and cell-type gene expression profiles in tomato plants colonized by an arbuscular mycorrhizal fungus. New Phytologist 184(4): 975–987.

Fiorilli V, Vannini C, Ortolani F, Garcia-Seco D, Chiapello M, Novero M, Domingo G, Terzi V, Morcia C, Bagnaresi P, et al. 2018. Omics approaches revealed how arbuscular mycorrhizal symbiosis enhances yield and resistance to leaf pathogen in wheat. Scientific Reports 8.

Floss DS, Levy JG, Levesque-Tremblay V, Pumplin N, Harrison MJ. 2013. DELLA proteins regulate arbuscule formation in arbuscular mycorrhizal symbiosis. Proceedings Of The National Academy Of Sciences Of The United States Of America 110(51): E5025–E5034.

Gao LL, Knogge W, Delp G, Smith FA, Smith SE. 2004. Expression patterns of defense-related genes in different types of arbuscular mycorrhizal development in wild-type and mycorrhiza-defective mutant tomato. Molecular Plant-Microbe Interactions 17(10): 1103–1113.

García-Garrido JM, Ocampo JA. 2002. Regulation of the plant defence response in arbuscular mycorrhizal symbiosis. Journal Of Experimental Botany 53(373): 1377–1386.

Gaude N, Schulze WX, P. F, Krajinski F. 2012. Cell type-specific protein and transcription profiles implicate periarbuscular membrane synthesis as an important carbon sink in the mycorrhizal symbiosis. Plant Signaling and Behavior 7(4): 461–464.

Gianinazzi-Pearson V. 1996. Plant cell responses to arbuscular mycorrhizal fungi: Getting to the roots of the symbiosis. Plant Cell 8(10): 1871–1883.

Gianinazzi-Pearson V, Dumas-Gaudot E, Gollotte A, Tahiri-Alaoui A, Gianinazzi S. 1996. Cellular and molecular defence-related root responses to invasion by arbuscular mycorrhizal fungi. New Phytologist 133(1): 45–57.

Gobbato E, Marsh JF, Vernie T, Wang E, Maillet F, Kim J, Miller JB, Sun J, Bano SA, Ratet P, et al. 2012. A GRAS-type transcription factor with a specific function in mycorrhizal signaling. Current Biology 22(23): 2236–2241.

Gollotte A, Gianinazzi-Pearson V, Giovannetti M, Sbrana C, Avio L, Gianinazzi S. 1993. Cellular localization and cytochemical probing of resistance reactions to arbuscular mycorrhizal fungi in a locus a myc-mutant of *Pisum sativum* L. Planta 191(1): 112–122.

Grunwald U, Nyamsuren O, Tamasloukht M, Lapopin L, Becker A, Mann P, Gianinazzi-Pearson V, Krajinski F, Franken P. 2004. Identification of mycorrhiza-regulated genes with arbuscule development-related expression profile. Plant Molecular Biology 55(4): 553–566.

Guenoune D, Galili S, Phillips DA, Volpin H, Chet I, Okon Y, Kapulnik Y. 2001. The defense response elicited by the pathogen *Rhizoctonia solani* is suppressed by colonization of the AM-fungus *Glomus intraradices*. Plant Science 160(5): 925–932.

Gutjahr C, Parniske M. 2013. Cell and developmental biology of arbuscular mycorrhiza symbiosis. Annual Review Of Cell And Developmental Biology 29: 593–617.

Handa Y, Nishide H, Takeda N, Suzuki Y, Kawaguchi M, Saito K. 2015. RNA-seq transcriptional profiling of an arbuscular mycorrhiza provides insights into regulated and coordinated gene expression in *Lotus japonicus* and *Rhizophagus irregularis*. Plant And Cell Physiology 56(8): 1490–1511.

Hückelhoven R 2007. Cell wall-associated mechanisms of disease resistance and susceptibility. Annual Review Of Phytopathology, 101–127.

Jones JDG, Dangl JL. 2006. The plant immune system. Nature 444(7117): 323–329.

Jung SC, Martinez-Medina A, Lopez-Raez JA, Pozo MJ. 2012. Mycorrhiza-induced resistance and priming of plant defenses. Journal Of Chemical Ecology 38(6): 651–664.

Kalo P, Gleason C, Edwards A, Marsh J, Mitra RM, Hirsch S, Jakab J, Sims S, Long SR, Rogers J, et al. 2005. Nodulation signaling in legumes requires NSP2, a member of the GRAS family of transcriptional regulators. Science 308(5729): 1786–1789.

Kamel L, Tang NW, Malbreil M, San Clemente H, Le Marquer M, Roux C, Frey NFD. 2017. The comparison of expressed candidate secreted proteins from two arbuscular mycorrhizal fungi unravels common and specific molecular tools to invade different host plants. Frontiers in Plant Science 8: 124.

Kapulnik Y, Volpin H, Itzhaki H, Ganon D, Galili S, David R, Shaul O, Elad Y, Chet I, Okon Y. 1996. Suppression of defence responses in mycorrhizal alfalfa and tobacco roots. New Phytologist 133(1): 59–64.

Klironomos JN. 2003. Variation in plant response to native and exotic arbuscular mycorrhizal fungi. Ecology 84(9): 2292–2301.

Kloppholz S, Kuhn H, Requena N. 2011. A secreted fungal effector of *Glomus intraradices* promotes symbiotic biotrophy. Current Biology 21(14): 1204–1209.

Kumar D, Yusuf MA, Singh P, Sardar M, Sarin NB. 2014. Histochemical detection of superoxide and H2O2 accumulation in *Brassica juncea* seedlings. Bio Protocols 4(8).

Kurihara D, Mizuta Y, Sato Y, Higashiyama T. 2015. ClearSee: a rapid optical clearing reagent for whole-plant fluorescence imaging. Development 142(23): 4168–4179.

Lanfranco L, Fiorilli V, Gutjahr C. 2018. Partner communication and role of nutrients in the arbuscular mycorrhizal symbiosis. New Phytologist 220(4): 1031–1046.

Lévy J, Bres C, Geurts R, Chalhoub B, Kulikova O, Duc G, Journet EP, Ané JM, Lauber E, Bisseling T, et al. 2004. A putative Ca2+ and calmodulin-dependent protein kinase required for bacterial and fungal symbioses. Science 303(5662): 1361–1364.

Liu CW, Breakspear A, Stacey N, Findlay K, Nakashima J, Ramakrishnan K, Liu MX, Xie F, Endre G, de Carvalho-Niebel F, et al. 2019. A protein complex required for polar growth of rhizobial infection threads. Nature Communications 10.

Liu JY, Maldonado-Mendoza I, Lopez-Meyer M, Cheung F, Town CD, Harrison MJ. 2007. Arbuscular mycorrhizal symbiosis is accompanied by local and systemic alterations in gene expression and an increase in disease resistance in the shoots. Plant Journal 50(3): 529–544.

Liu QQ, Luo L, Zheng LQ. 2018. Lignins: Biosynthesis and biological functions in plants. International Journal of Molecular Sciences 19(2).

Loake G, Grant M. 2007. Salicylic acid in plant defence-the players and protagonists. Current Opinion In Plant Biology 10(5): 466–472.

Maillet F, Poinsot V, André O, Puech-Pagès V, Haouy A, Gueunier M, Cromer L, Giraudet D, Formey D, Niebel A, et al. 2011. Fungal lipochitooligosaccharide symbiotic signals in arbuscular mycorrhiza. Nature 469: 58–64.

Marcel S, Sawers R, Oakeley E, Angliker H, Paszkowski U. 2010. Tissue-adapted invasion strategies of the rice blast fungus *Magnaporthe oryzae*. Plant Cell 22(9): 3177–3187.

Marsh JF, Schultze M. 2001. Analysis of arbuscular mycorrhizas using symbiosis-defective plant mutants. New Phytologist 150(3): 525–532.

Miedes E, Van Holme R, Boerjan W, Molina A. 2014. The role of the secondary cell wall in plant resistance to pathogens. Frontiers in Plant Science 5: 358.

Millet YA, Danna CH, Clay NK, Songnuan W, Simon MD, Werck-Reichhart D, Ausubel FM. 2010. Innate immune responses activated in Arabidopsis roots by microbe-associated molecular patterns. Plant Cell 22(3): 973–990.

Mitra RM, Gleason CA, Edwards A, Hadfield J, Downie JA, Oldroyd GED, Long SR. 2004. A Ca2+/calmodulin-dependent protein kinase required for symbiotic nodule development: Gene identification by transcript-based cloning. Proceedings Of The National Academy Of Sciences Of The United States Of America 101(13): 4701–4705.

Morin E, Miyauchi S, San Clemente H, Chen ECH, Pelin A, de la Providencia I, Ndikumana S, Beaudet D, Hainaut M, Drula E, et al. 2019. Comparative genomics of *Rhizophagus irregularis, R. cerebriforme, R. diaphanus* and *Gigaspora rosea* highlights specific genetic features in Glomeromycotina. New Phytologist 222(3): 1584–1598.

Murray JD, Duvvuru Muni R, Torres-Jerez I, Tang Y, Allen S, Andriankaja M, Li G, Laxmi A, Cheng X, Wen J, et al. 2011. *Vapyrin*, a gene essential for intracellular progression of arbuscular mycorrhizal symbiosis, is also essential for infection by rhizobia in the nodule symbiosis of *Medicago truncatula*. Plant Journal 65(2): 244–252.

Nouri E, Breuillin-Sessoms F, Feller U, Reinhardt D. 2014. Phosphorus and nitrogen regulate arbuscular mycorrhizal symbiosis in *Petunia hybrida*. Plos One 9(3): e90841.

Novero M, Faccio A, Genre A, Stougaard J, Webb KJ, Mulder L, Parniske M, Bonfante P. 2002. Dual requirement of the *LjSym4* gene for mycorrhizal development in epidermal and cortical cells of *Lotus japonicus* roots. New Phytologist 154(3): 741–749.

Park H-J, Floss DS, Levesque-Tremblay V, Bravo A, Harrison MJ. 2015. Hyphal branching during arbuscule development requires *Reduced Arbuscular Mycorrhiza1*. Plant Physiology 169(4): 2774–2788.

Pfaffl MW. 2001. A new mathematical model for relative quantification in real-time RT-PCR. Nucleic Acids Research 29(9).

Pieterse CMJ, Zamioudis C, Berendsen RL, Weller DM, Van Wees SCM, Bakker P 2014. Induced systemic resistance by beneficial microbes. In: VanAlfen NK ed. Annual Review of Phytopathology, Vol 52, 347–375.

Pimprikar P, Carbonnel S, Paries M, Katzer K, Klingl V, Bohmer MJ, Karl L, Floss DS, Harrison MJ, Parniske M, et al. 2016. A CCaMK-CYCLOPS-DELLA complex activates transcription of RAM1 to regulate arbuscule branching. Current Biology 26: 987–998.

Pozo MJ, Azcon-Aguilar C. 2007. Unraveling mycorrhiza-induced resistance. Current Opinion In Plant Biology 10(4): 393–398.

Pumplin N, Mondo SJ, Topp S, Starker CG, Gantt JS, Harrison MJ. 2010. *Medicago truncatula* Vapyrin is a novel protein required for arbuscular mycorrhizal symbiosis. Plant Journal 61(3): 482–494.

Reynolds ES. 1963. The use of lead citrate at high pH as an electron-opaque stain in electron microscopy. Journal Of Cell Biology 17: 208–212.

Rich MK, Courty P-E, Roux C, Reinhardt D. 2017. Role of the GRAS transcription factor ATA/RAM1 in the transcriptional reprogramming of arbuscular mycorrhiza in *Petunia hybrida*. Bmc Genomics 18: 589.

Rich MK, Schorderet M, Bapaume L, Falquet L, Morel P, Vandenbussche M, Reinhardt D. 2015. A petunia GRAS transcription factor controls symbiotic gene expression and fungal morphogenesis in arbuscular mycorrhiza. Plant Physiol 168: 788–797.

Rich MK, Schorderet M, Reinhardt D. 2014. The role of the cell wall compartment in mutualistic symbioses of plants. Frontiers in Plant Science 5: 238.

Ruiz-Lozano JM, Roussel H, Gianinazzi S, Gianinazzi-Pearson V. 1999. Defense genes are differentially induced by a mycorrhizal fungus and Rhizobium sp in wild-type and symbiosis-defective pea genotypes. Molecular Plant-Microbe Interactions 12(11): 976–984.

Salzer P, Bonanomi A, Beyer K, Vögeli-Lange R, Aeschbacher RA, Lange J, Wiemken A, Kim D, Cook DR, Boller T. 2000. Differential expression of eight chitinase genes in *Medicago truncatula* roots during mycorrhiza formation, nodulation, and pathogen infection. Molecular Plant-Microbe Interactions 13(7): 763–777.

Salzer P, Corbière H, Boller T. 1999. Hydrogen peroxide accumulation in *Medicago truncatula* roots colonized by the arbuscular mycorrhiza-forming fungus *Glomus intraradices*. Planta 208(3): 319–325.

Scheler C, Durner J, Astier J. 2013. Nitric oxide and reactive oxygen species in plant biotic interactions. Current Opinion In Plant Biology 16(4): 534–539.

Sedzielewska Toro K, Brachmann A. 2016. The effector candidate repertoire of the arbuscular mycorrhizal fungus *Rhizophagus clarus*. Bmc Genomics 17: 101.

Sekhara Reddy DMR, Schorderet M, Feller U, Reinhardt D. 2007. A petunia mutant affected in intracellular accommodation and morphogenesis of arbuscular mycorrhizal fungi. Plant Journal 51: 739–750.

Siciliano V, Genre A, Balestrini R, Cappellazzo G, deWit P, Bonfante P. 2007. Transcriptome analysis of arbuscular mycorrhizal roots during development of the prepenetration apparatus. Plant Physiology 144(3): 1455–1466.

Smith FA, Grace EJ, Smith SE. 2009. More than a carbon economy: nutrient trade and ecological sustainability in facultative arbuscular mycorrhizal symbioses. New Phytologist 182(2): 347–358.

Smith SE, Read DJ. 2008. Mycorrhizal Symbiosis. New York: Academic Press.

Spatafora JW, Chang Y, Benny GL, Lazarus K, Smith ME, Berbee ML, Bonito G, Corradi N, Grigoriev I, Gryganskyi A, et al. 2016. A phylum-level phylogenetic classification of zygomycete fungi based on genome-scale data. Mycologia 108(5): 1028–1046.

Spurr AR. 1969. A low-viscosity epoxy resin embedding medium for electron microscopy. Journal of Ultrastructure Research 26(1-2): 31–43.

Stuurman J, Kuhlemeier C. 2005. Stable two-element control of *dTph1* transposition in mutator strains of *Petunia* by an inactive *ACT1* introgression from a wild species. Plant Journal 41(6): 945–955.

Tang N, San Clemente H, Roy S, Bécard G, Zhao B, Roux C. 2016. A survey of the gene repertoire of *Gigaspora rosea* unravels conserved features among Glomeromycota for obligate biotrophy. Frontiers in Microbiology 7: 233.

Thuerig B, Felix G, Binder A, Boller T, Tamm L. 2005. An extract of *Penicillium chrysogenum* elicits early defense-related responses and induces resistance in *Arabidopsis thaliana* independently of known signalling pathways. Physiological And Molecular Plant Pathology 67(3-5): 180–193.

Torres MA, Jones JDG, Dangl JL. 2006. Reactive oxygen species signaling in response to pathogens. Plant Physiology 141(2): 373–378.

van Loon LC, Geraats BPJ, Linthorst HJM. 2006a. Ethylene as a modulator of disease resistance in plants. Trends In Plant Science 11(4): 184–191.

van Loon LC, Rep M, Pieterse CMJ. 2006b. Significance of inducible defense-related proteins in infected plants. Annual Review Of Phytopathology 44: 135–162.

Vanholme R, De Meester B, Ralph J, Boerjan W. 2019. Lignin biosynthesis and its integration into metabolism. Current Opinion In Biotechnology 56: 230–239.

Vierheilig H, Alt M, Lange J, Gutrella M, Wiemken A, Boller T. 1995. Colonization of transgenic tobacco constitutively expressing pathogenesis-related proteins by the vesicular-arbuscular mycorrhizal fungus *Glomus mosseae*. Applied And Environmental Microbiology 61(8): 3031–3034.

Vlot AC, Dempsey DA, Klessig DF 2009. Salicylic acid, a multifaceted hormone to combat disease. Annual Review Of Phytopathology, 177–206.

Vogt T. 2010. Phenylpropanoid biosynthesis. Molecular Plant 3(1): 2–20.

Voss S, Betz R, Heidt S, Corradi N, Requena N. 2018. RiCRN1, a crinkler effector from the arbuscular mycorrhizal fungus *Rhizophagus irregularis*, functions in arbuscule development. Frontiers in Microbiology 9: 2068.

Wasternack C, Hause B. 2013. Jasmonates: biosynthesis, perception, signal transduction and action in plant stress response, growth and development. An update to the 2007 review in Annals of Botany. Annals Of Botany 111(6): 1021–1058.

Wegel E, Schauser L, Sandal N, Stougaard J, Parniske M. 1998. Mycorrhiza mutants of Lotus japonicus define genetically independent steps during symbiotic infection. Molecular Plant-Microbe Interactions 11(9): 933–936.

Zhang XC, Pumplin N, Ivanov S, Harrison MJ. 2015. EXO70I is required for development of a sub-domain of the periarbuscular membrane during arbuscular mycorrhizal symbiosis. Current Biology 25(16): 2189–2195.

Zipfel C, Oldroyd GED. 2017. Plant signalling in symbiosis and immunity. Nature 543(7645): 328–336.

