## Supplemental Information for "VAPYRIN attenuates defence by repressing PR gene induction and localized lignin accumulation during arbuscular mycorrhizal symbiosis of *Petunia hybrida*"

### Chen et al. – Supplementary Information

#### Figure S1

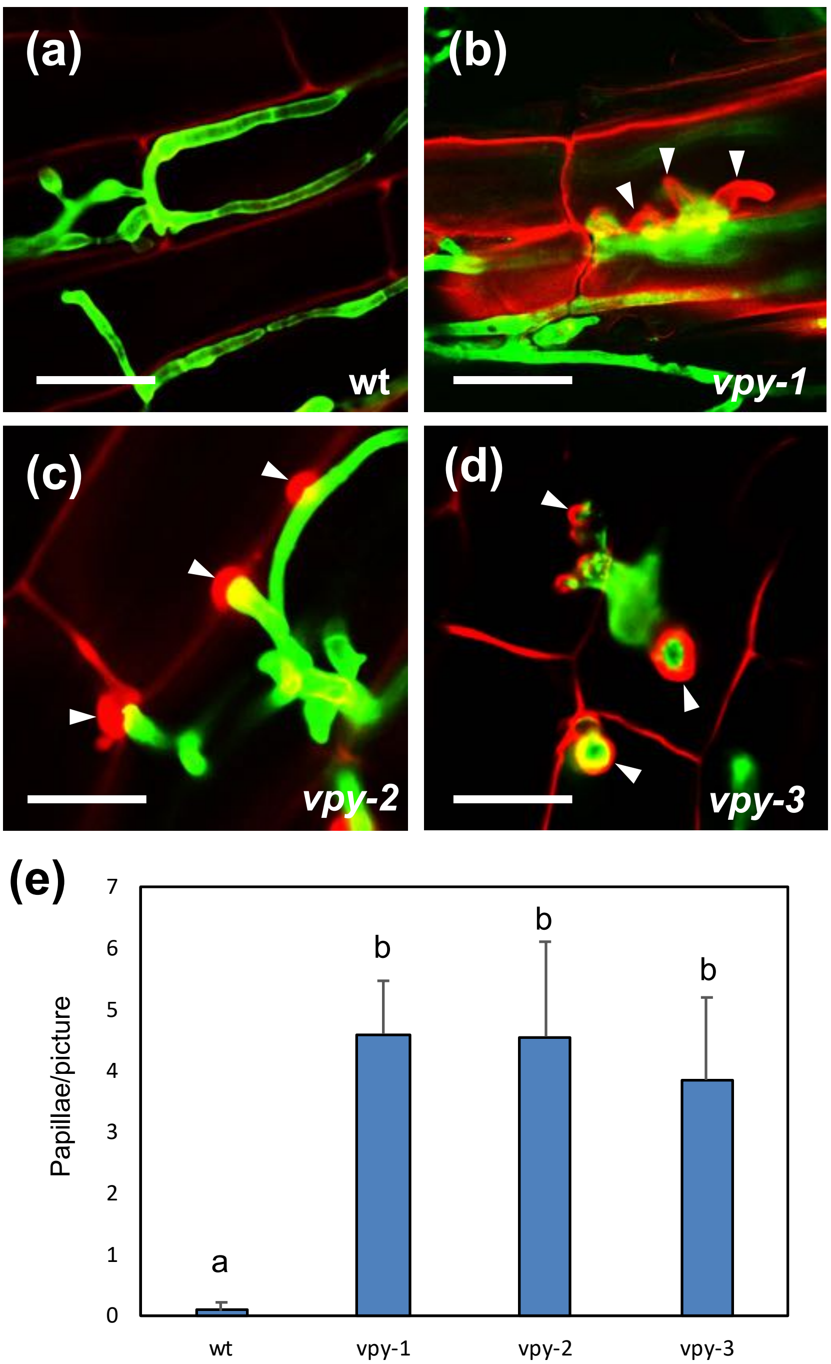

Figure S2

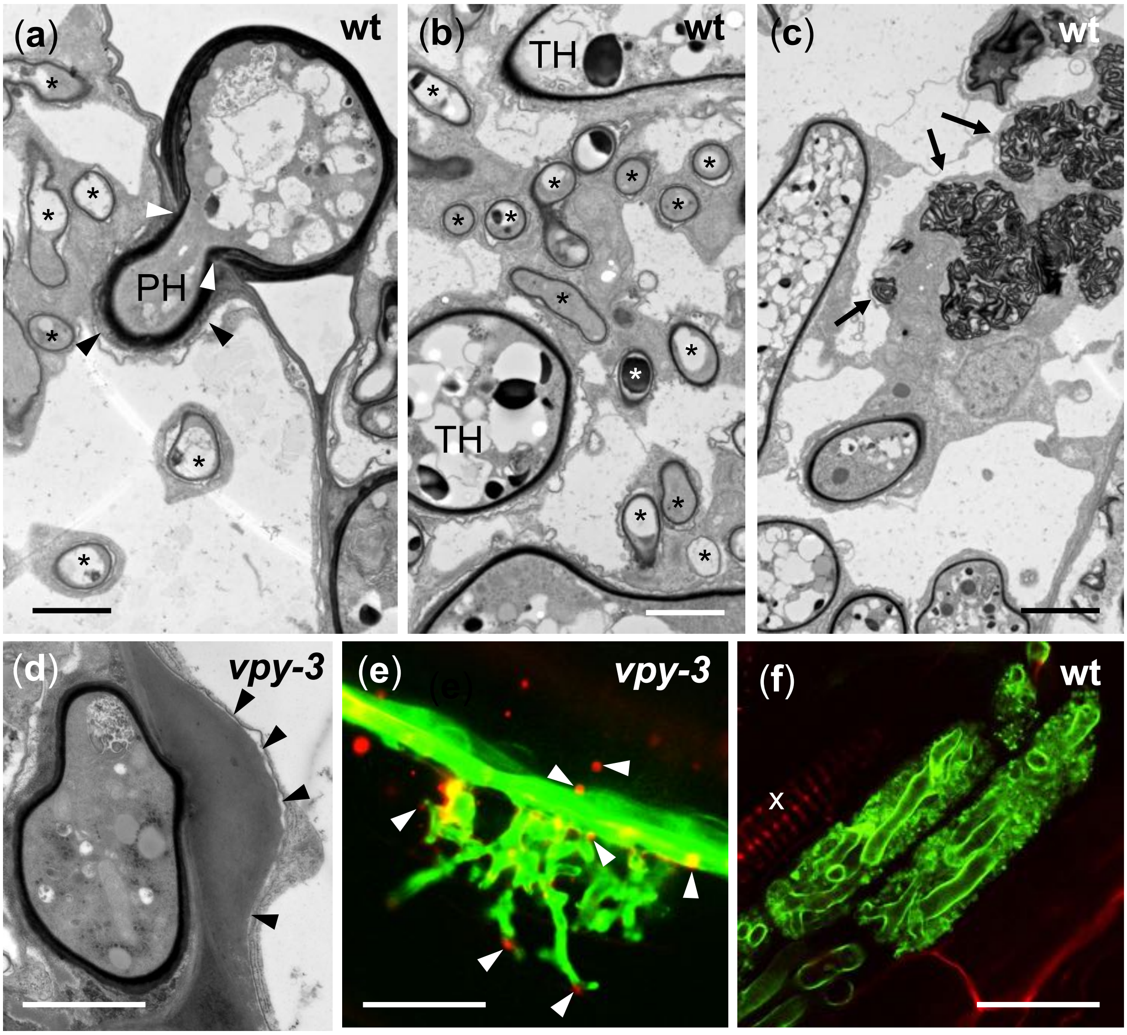

Figure S3

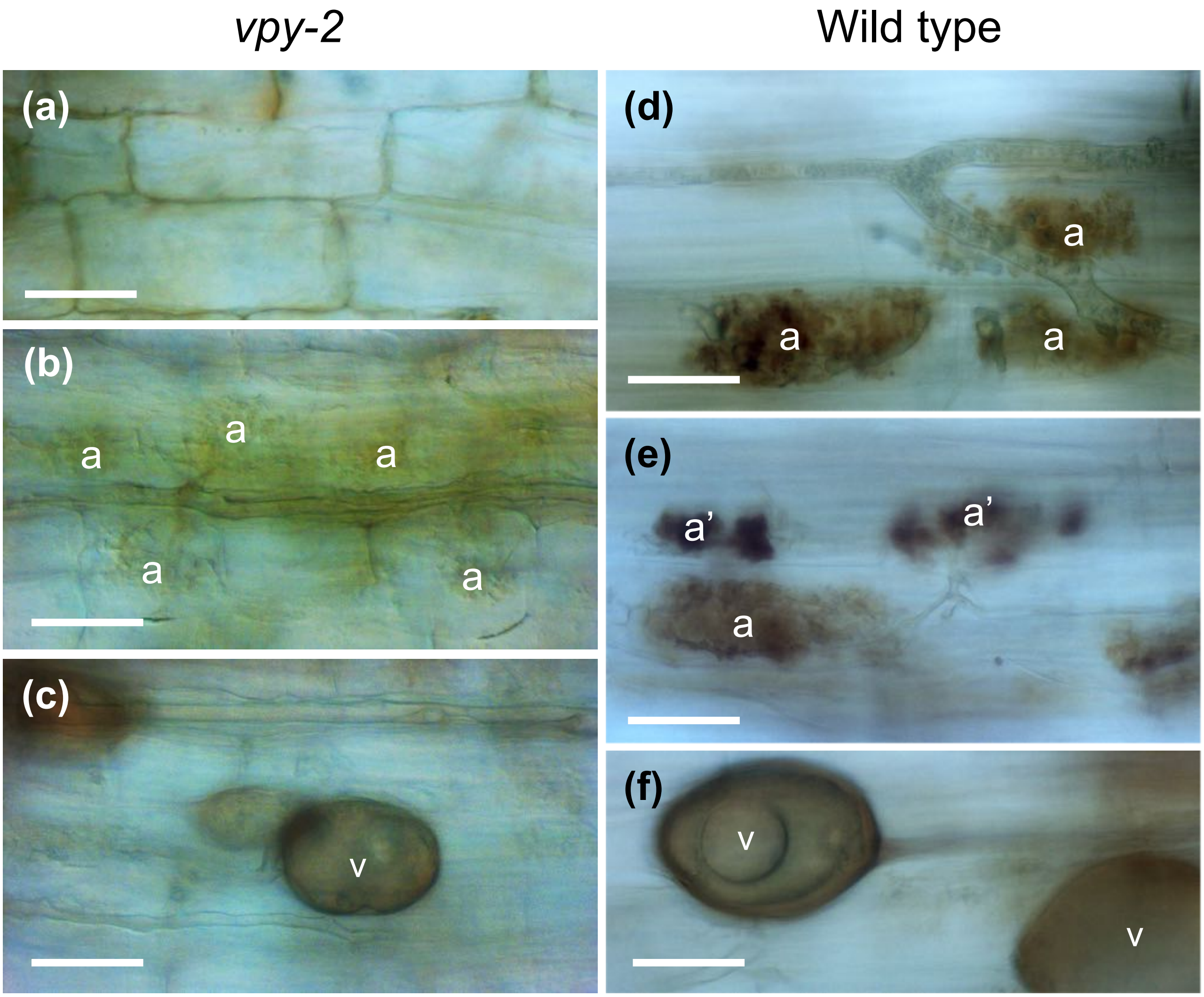

Figure S4

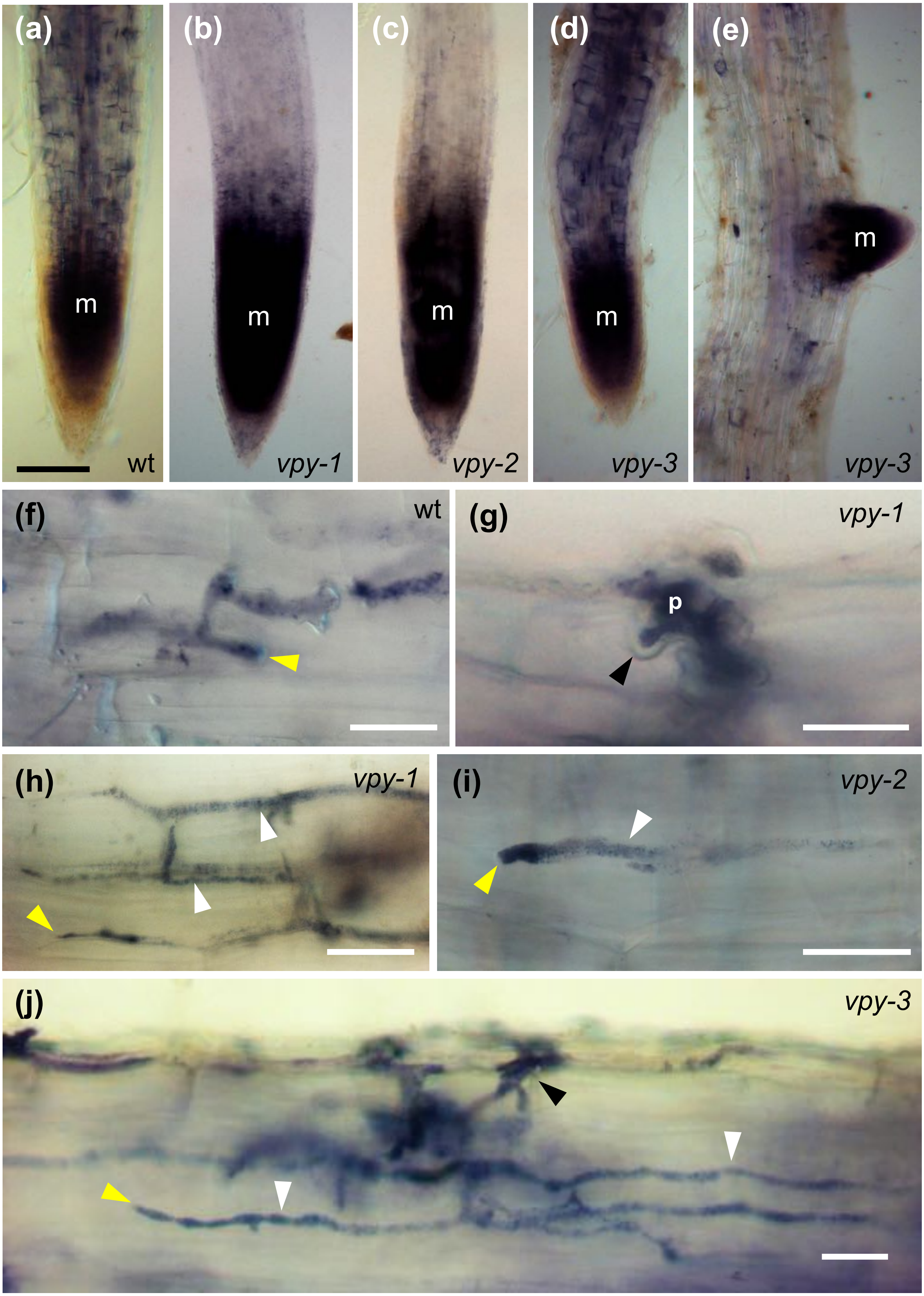

(a)
m
wt
(b)
m
vpy-1
(c)
m
vpy-2
(d)
m
vpy-3
(e)
m
vpy-3
(f)
wt
(g)
vpy-1
p
(h)
vpy-1
(i)
vpy-2
(j)
vpy-3

Figure S5

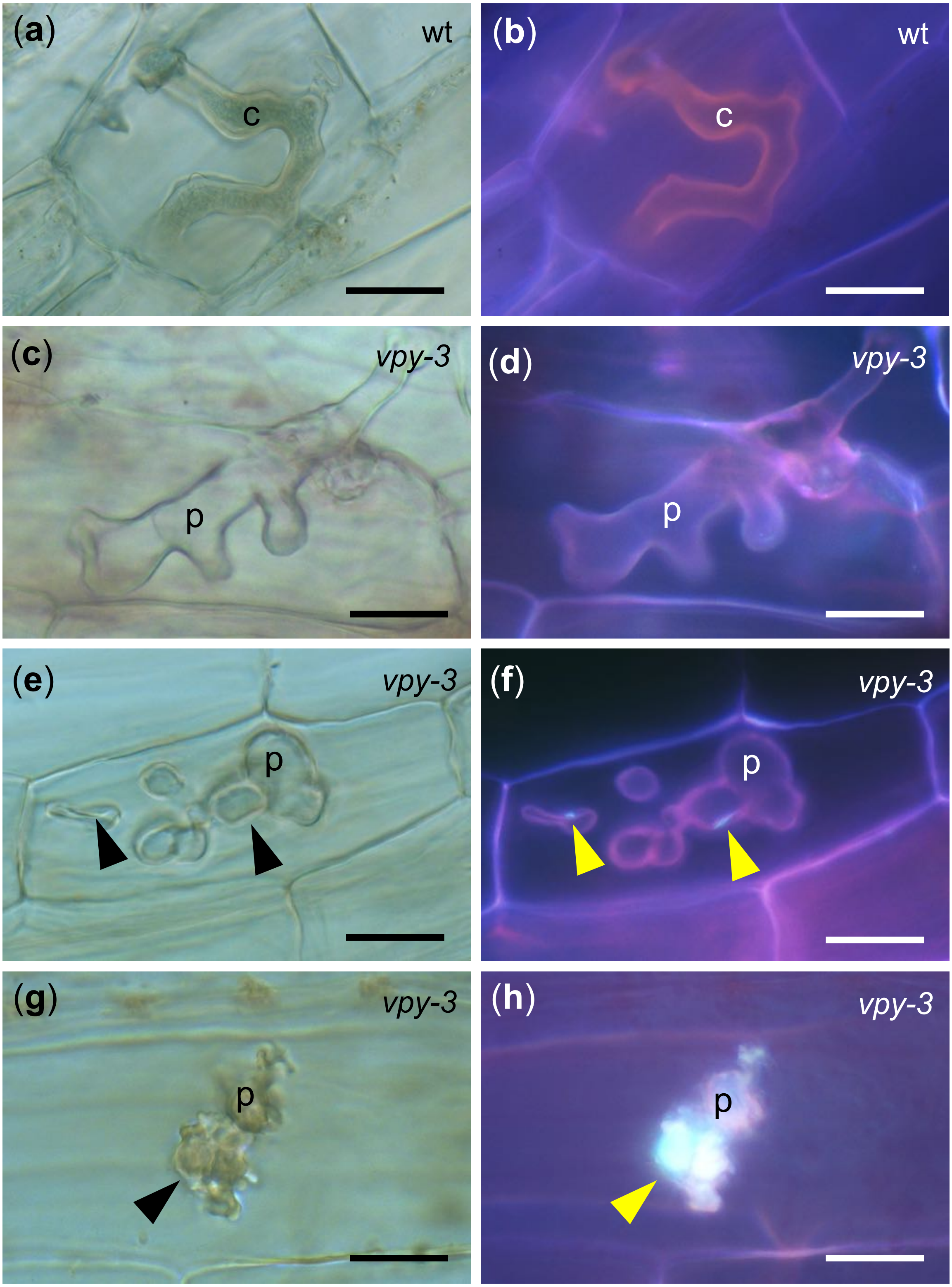

Figure S6

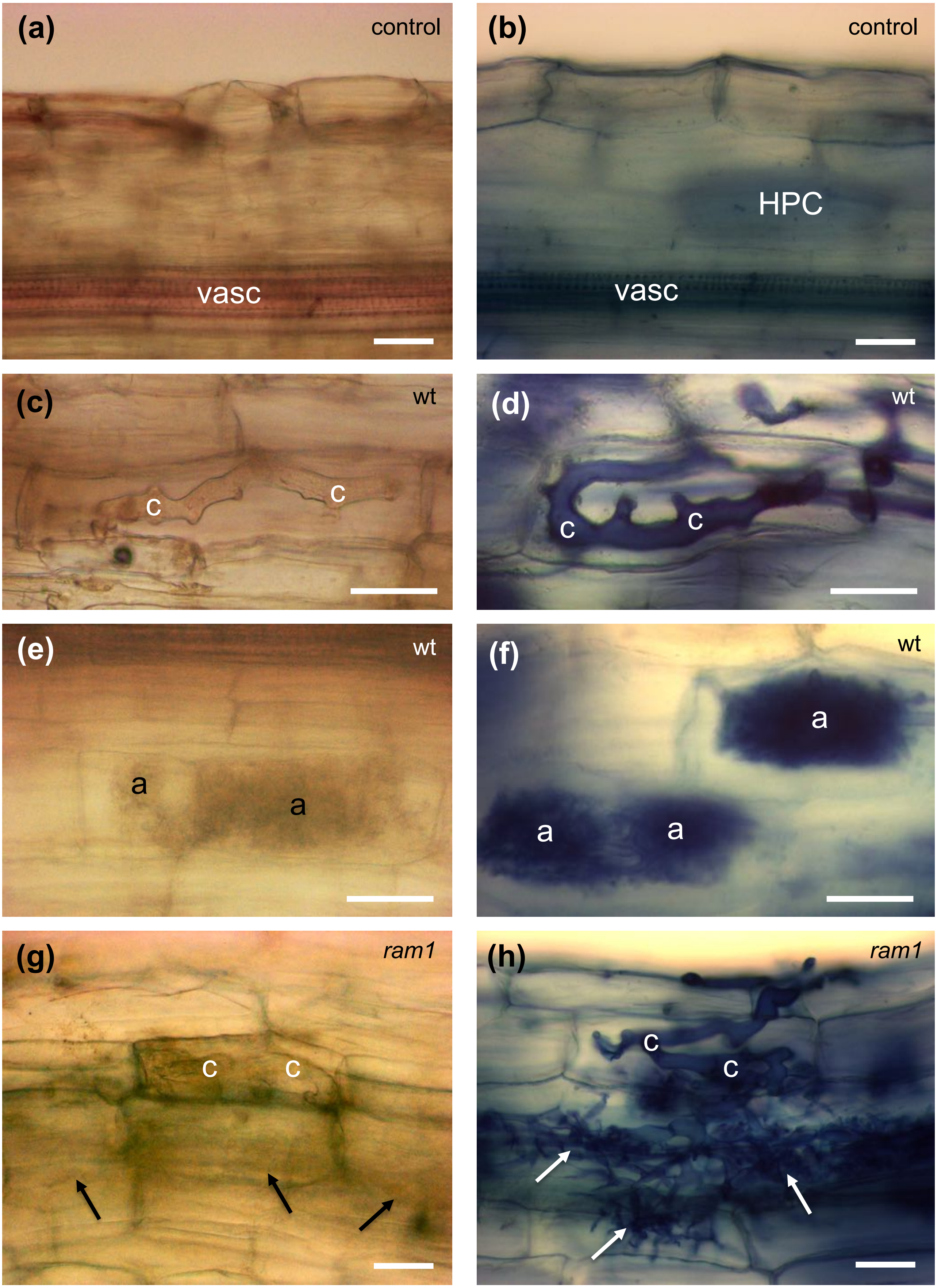
(a)
control
vasc
(b)
control
HPC
vasc
(c)
wt
c
c
(d)
wt
c
c
(e)
wt
a
a
(f)
wt
a
a
a
(g)
ram1
c
c
(h)
ram1
c
c

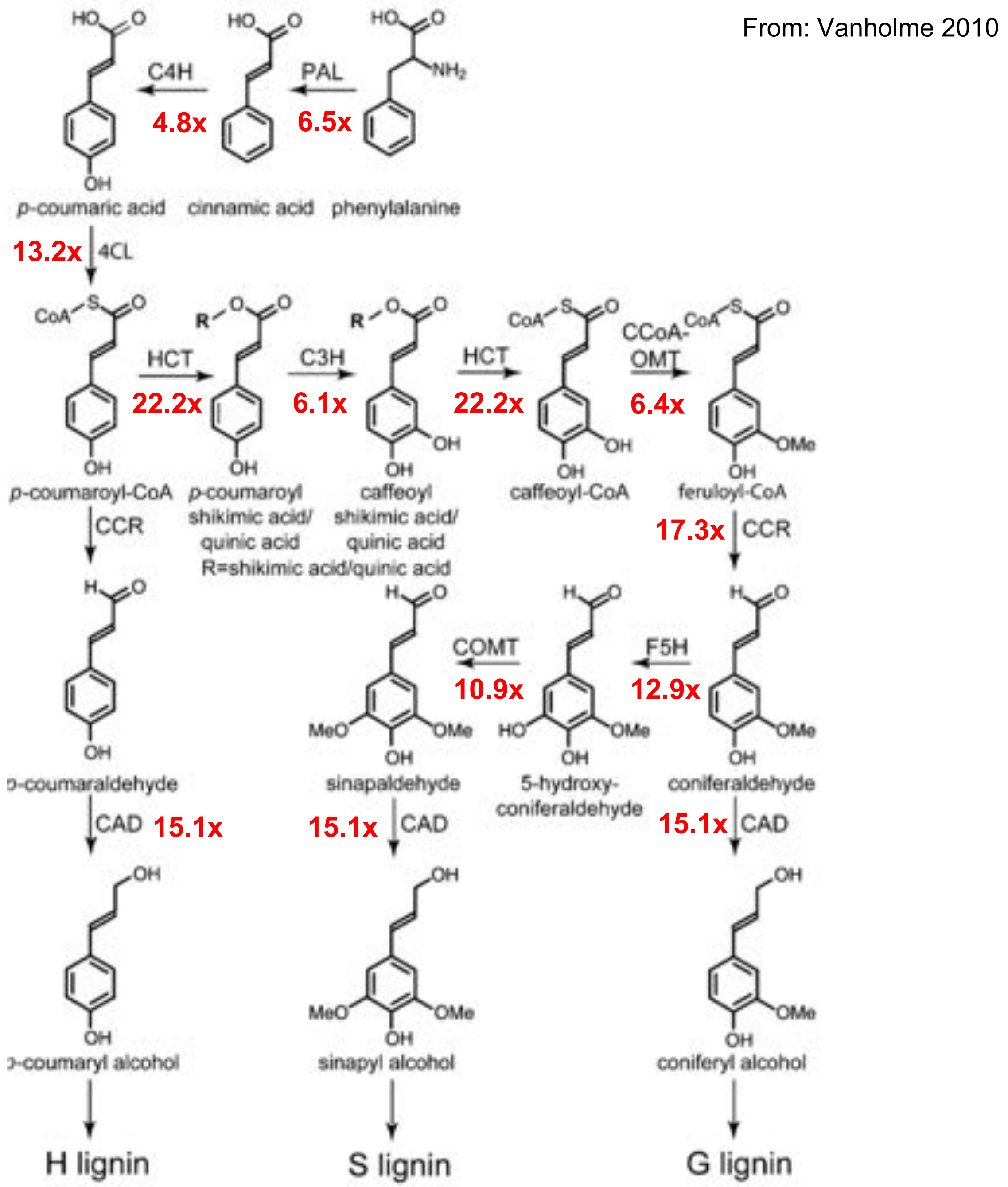


### Figure S7. Activation of the lignin biosynthetic pathway in mycorrhizal *vpy* mutants

Figure 1. The main biosynthetic route toward the monolignols p-coumaryl, coniferyl, and sinapyl alcohol (Boerjan et al., 2003).
PAL, PHENYLALANINE AMMONIA-LYASE;
C4H, CINNAMATE 4-HYDROXYLASE;
4CL, 4-COUMARATE:CoA LIGASE;
C3H, p-COUMARATE 3-HYDROXYLASE;
HCT, p-HYDROXYCINNAMOYL-CoA:QUINATE/ SHIKIMATE p-HYDROXYCINNAMOYLTRANSFERASE;
CCoAOMT, CAFFEOYL-CoA O-METHYLTRANSFERASE;
CCR, CINNAMOYL-CoA REDUCTASE;
F5H, FERULATE 5-HYDROXYLASE;
COMT, CAFFEIC ACID O-METHYLTRANSFERASE;
CAD, CINNAMYL ALCOHOL DEHYDROGENASE.

### Figure S8

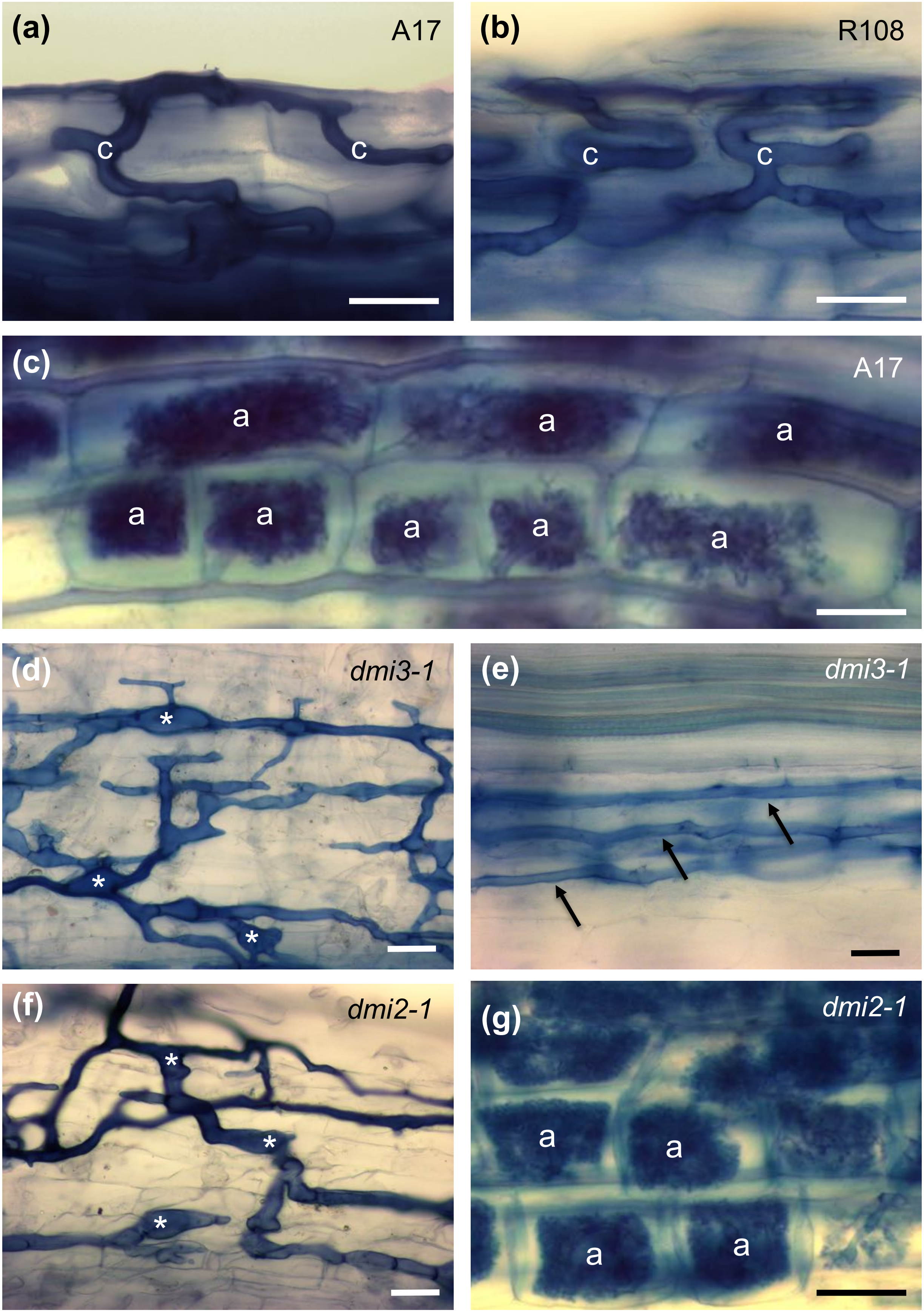

(a)
A17
c
c
(b)
R108
c
c
(c)
A17
a
a
a
a
a
a
a
a
(d)
dmi3-1
*
*
*
(e)
dmi3-1
(f)
dmi2-1
*
*
*
(g)
dmi2-1
a
a
a
a

Figure S9

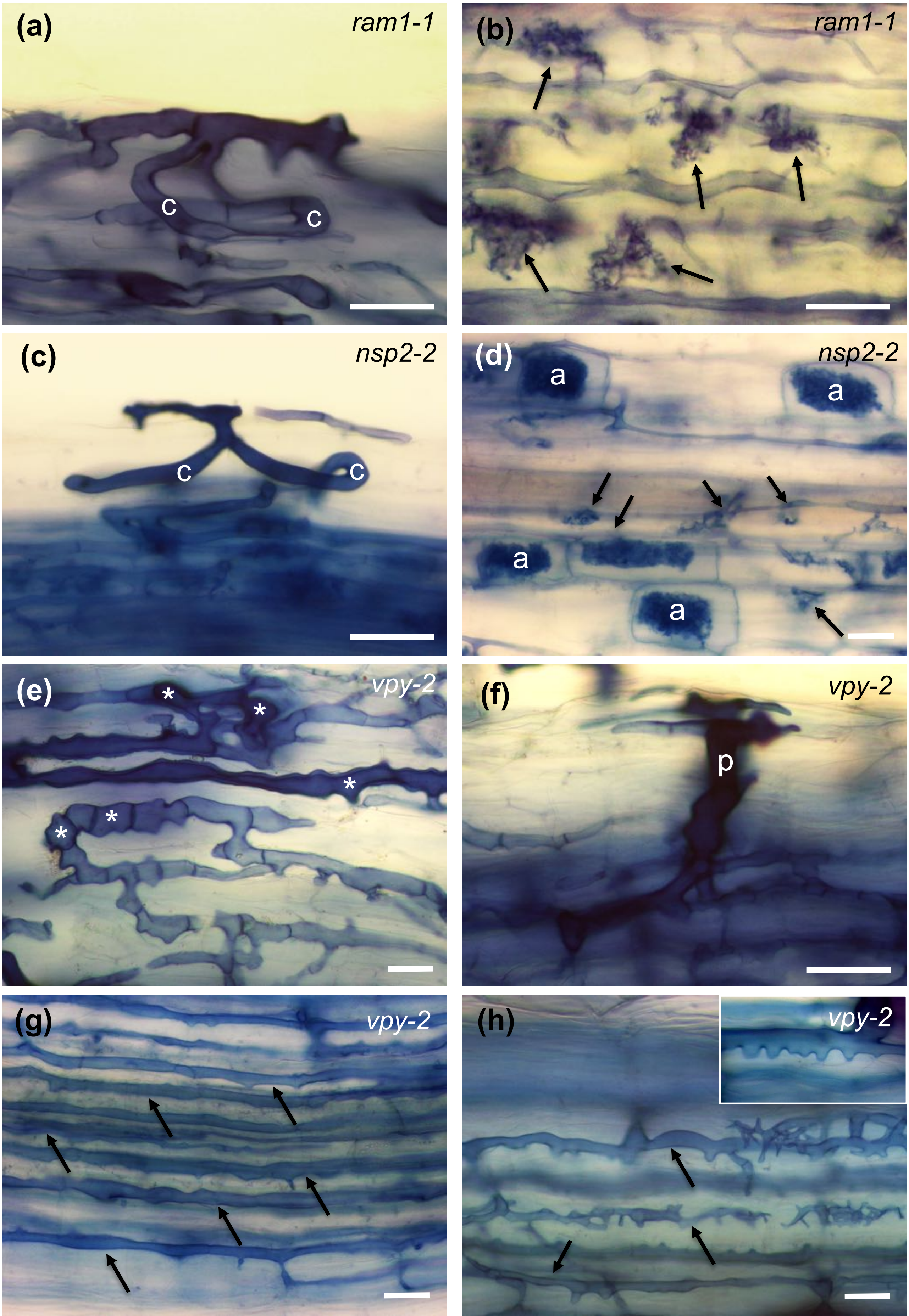
(a)
ram1-1
c
c
(b)
ram1-1
(c)
nsp2-2
c
c
(d)
nsp2-2
a
a
a
a
(e)
vpy-2
*
*
*
*
*
(f)
vpy-2
p
(g)
vpy-2
(h)
vpy-2

Figure S10

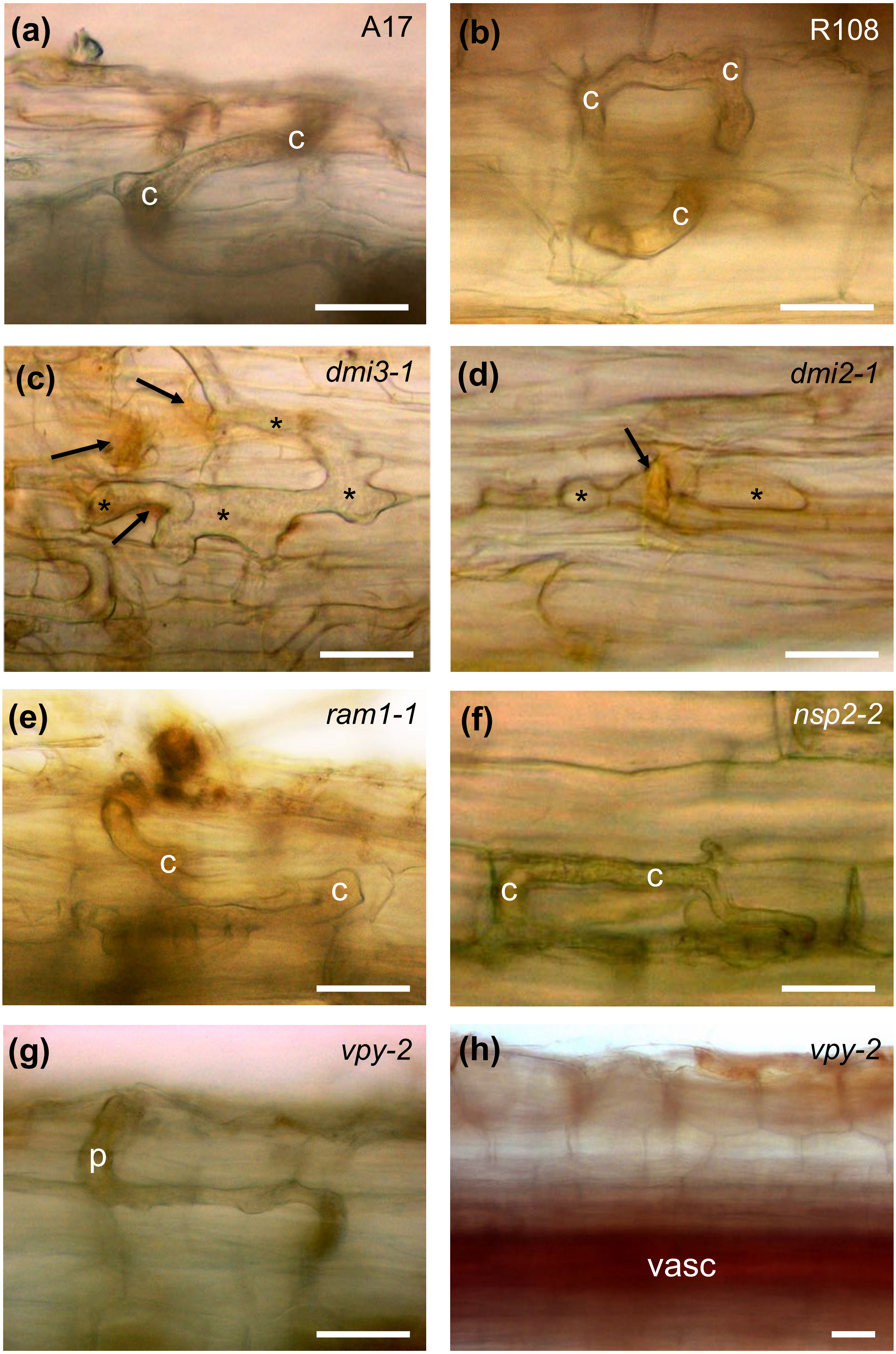

(a)
A17
c
c
(b)
R108
c
c
c
(c)
*dmi3-1*
*
*
*
*
(d)
*dmi2-1*
*
*
(e)
*ram1-1*
c
c
(f)
*nsp2-2*
c
c
(g)
*vpy-2*
p
(h)
*vpy-2*
vasc

### Figure S11. SA levels

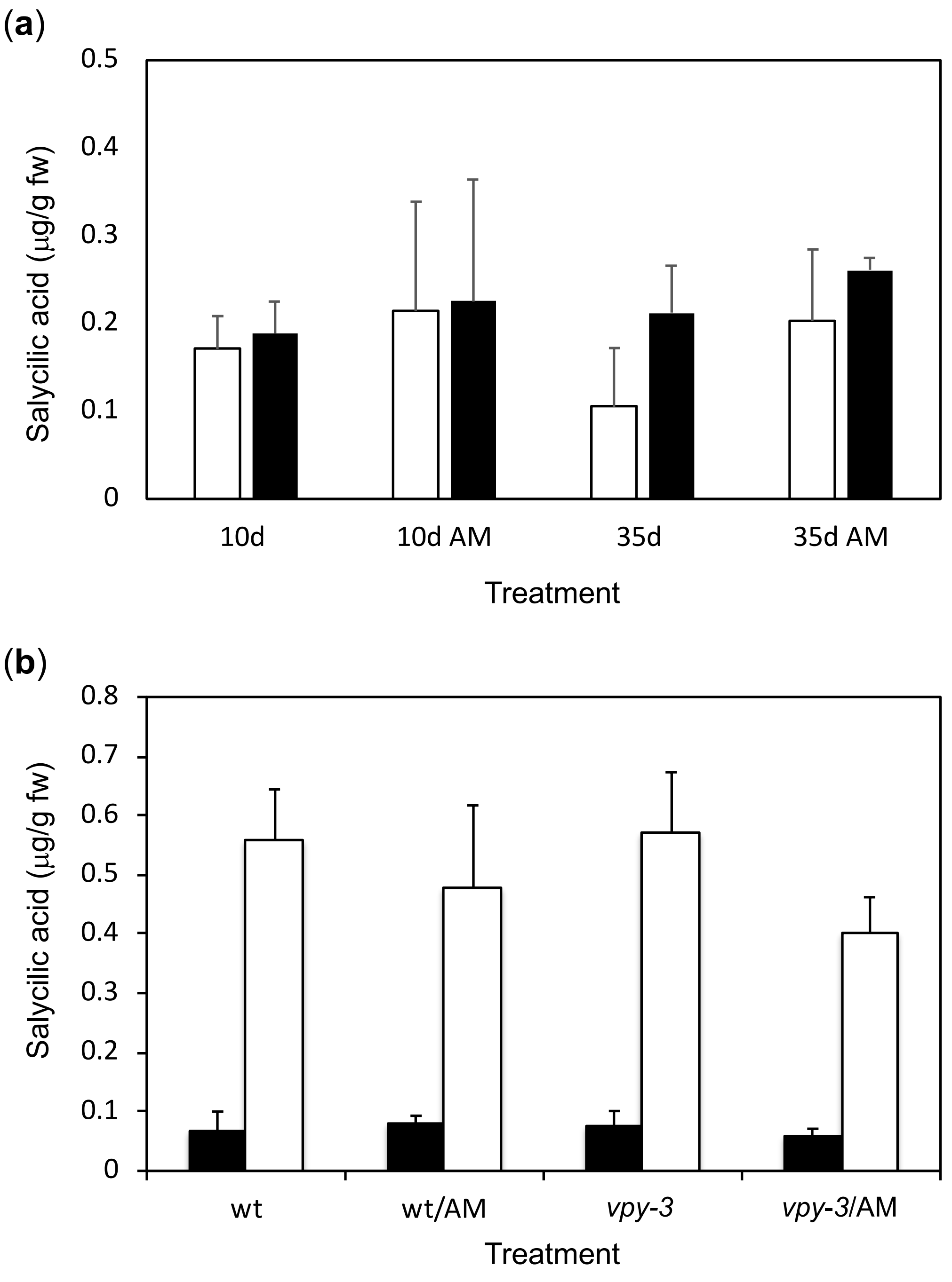

Figure S12. JA levels

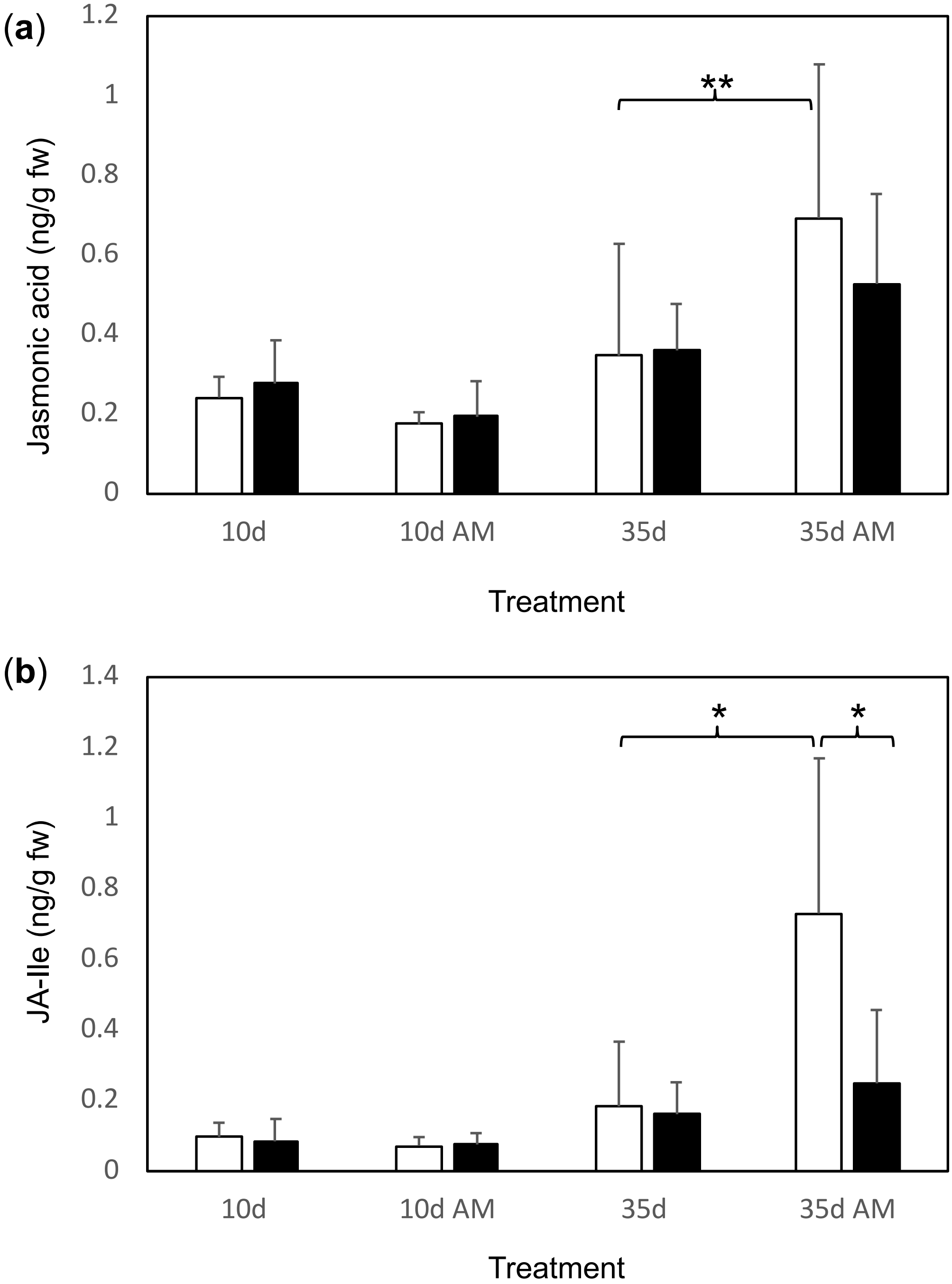

### Figure S13

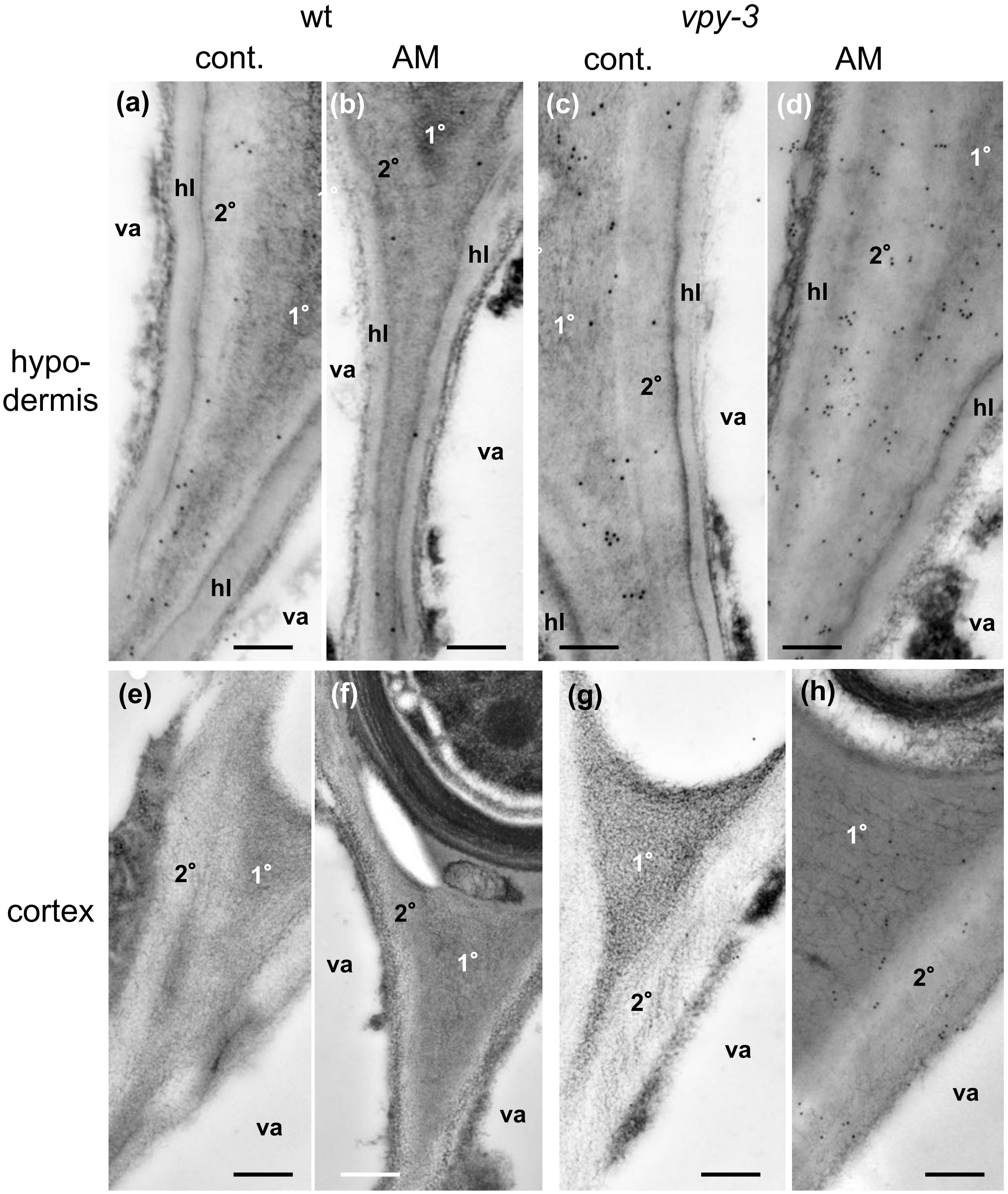

Figure S14

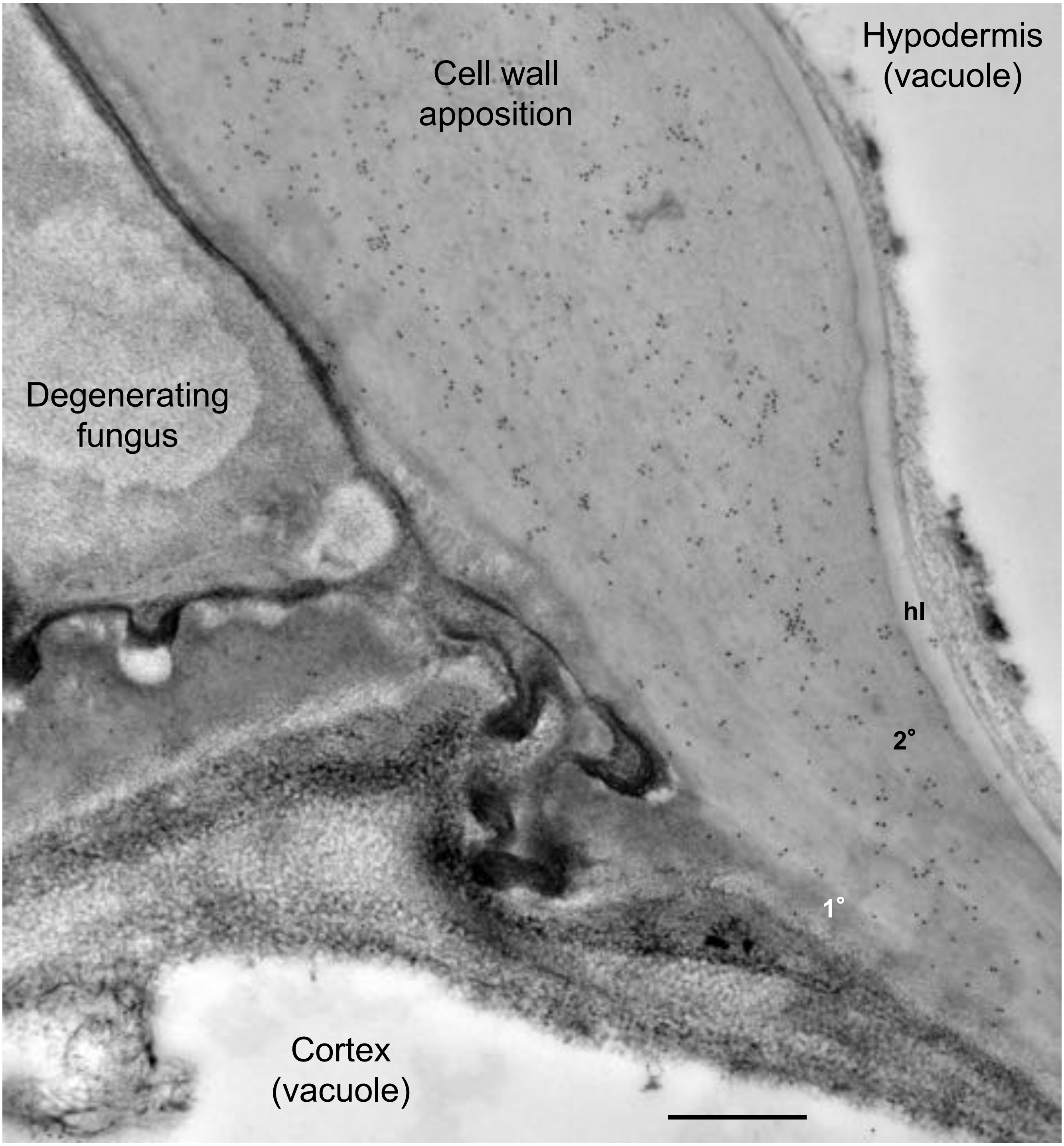

Cell wall
apposition
Hypodermis
(vacuole)
Degenerating
fungus
hl
2°
1°
Cortex
(vacuole)

Figure S15.

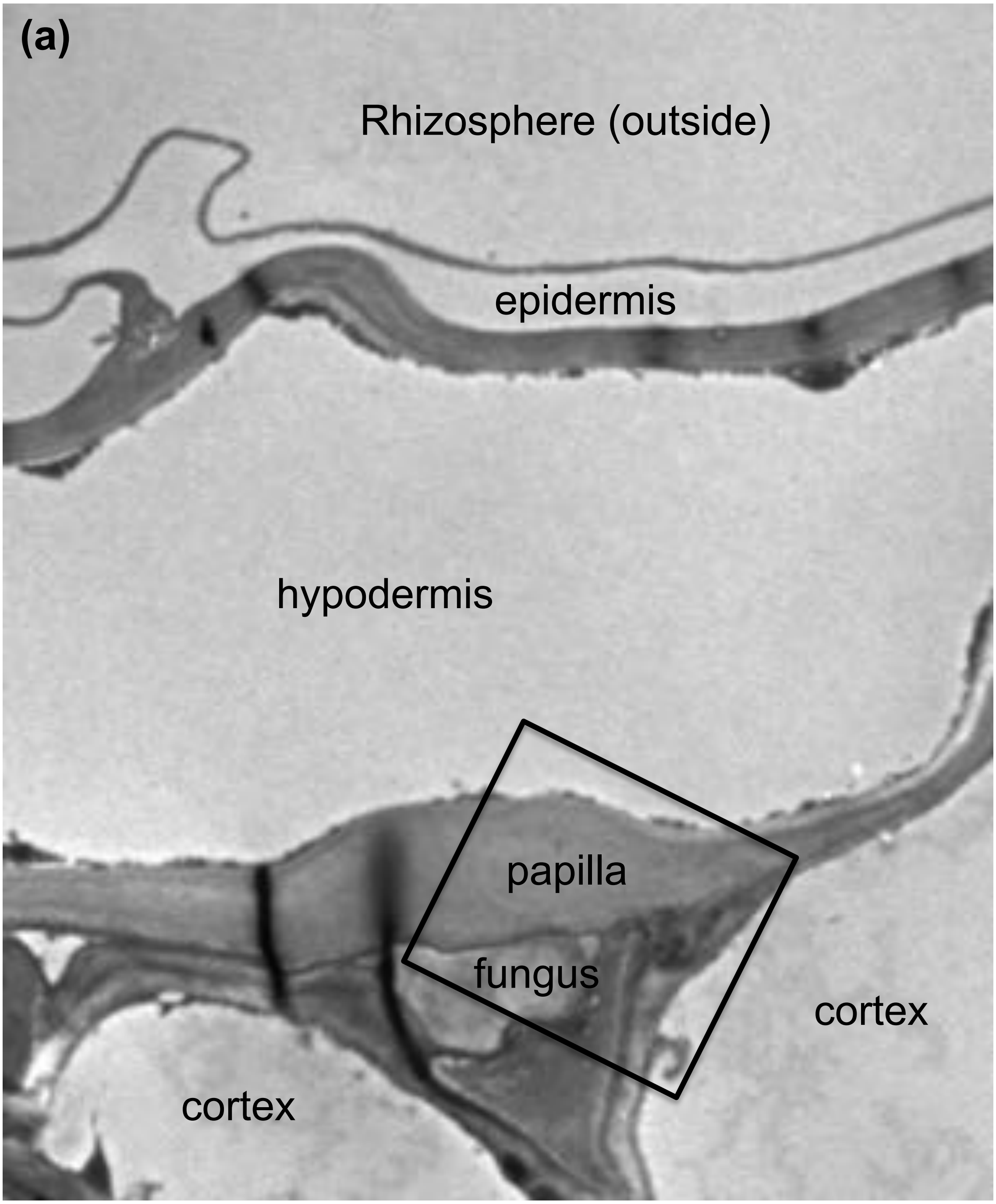
(a)
Rhizosphere (outside)
epidermis
hypodermis
papilla
fungus
cortex
cortex

### Figure S16. Quantification of gold particles

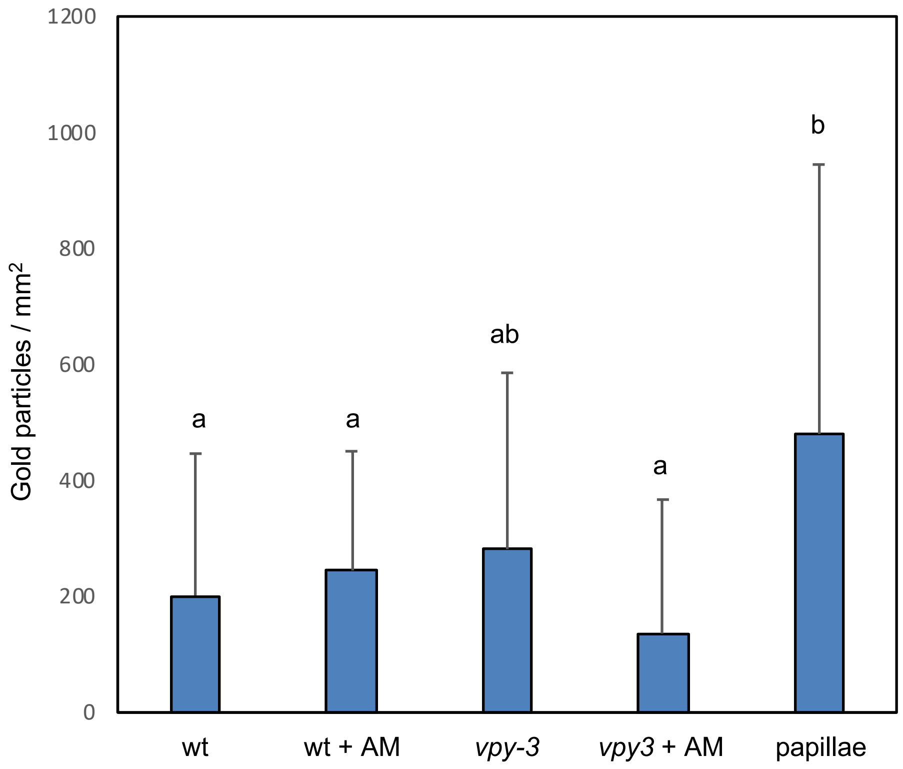

### Table S1 - Callose

| | | | Callose signal | | | | | |
|---|---|---|---|---|---|---|---|---|
| | Evaluated sites | infected cells | strong | weak | negative | | callose all | % callose |
| wt | 30 | 73 | 0 | 3 | 70 | | 3 | 4.11 |
| *vpy-1* | 38 | 113 | 0 | 2 | 111 | | 2 | 1.77 |
| *vpy-2* | 40 | 79 | 2 | 6 | 68 | | 8 | 10.13 |
| *vpy-3* | 38 | 89 | 1 | 4 | 83 | | 5 | 5.62 |

Table S2

#### *M. truncatula* mutants

| Mutant | Affected gene | Genetic background | Reference |
| --- | --- | --- | --- |
| *dmi2-1* | *SYMRK* | A17 | Endre et al., 2002 |
| *dmi3-1* | *CCAMK* | A17 | Lévy et al., 2004; Mitra et al., 2004 |
| *ram1-1* | *RAM1* | A17 | Gobbato et al., 2012 |
| *nsp2-2* | *NSP* | A17 | Kalo et al., 2005 |
| *vpy-2* (NF6898.86) | *VYPYRIN* | R108 | Liu et al., 2019 |

### Table S3 – qPCR primers for lignin biosynthetic genes

| | Gene ID | Forward Primer | Reverse Primer |
|---|---|---|---|
| PAL1 | Peaxi162Scf00123g00096.1 | CAATGGGTGCTAATGGTGAACTTCAT | CCTTCAATTCATCTTCGAAAGCTCCA |
| PAL2 | Peaxi162Scf00488g00074.1 | AAGATTGCAGCTTTTGAAGACGAGTT | ACCGTTCCATTCCTTGAGACATTCTA |
| PAL3 | Peaxi162Scf00858g00215.1 | CTAGTGGCCAAGAAAGTGTTGACAAT | AGCTCTCTCAATTTCCTTAGGCAGAA |
| 4CL1 | Peaxi162Scf00314g00086.1 | TCCTAACATTTCTGATGCTGCTGTTG | GCAAGTCTAGCTCTCAAGTCCTTTCT |
| 4CL2 | Peaxi162Scf00207g00334.1 | GATCCTGATACTGGGTGGCTACATAC | TTCTCCTGCTTGTTCATCTTTCATGG |
| 4CL3 | Peaxi162Scf00745g00865.1 | GAGTCTCTTCTTGTAAGCCATCCAGA | CTGCTAGTTTGGCTCTCAGTTCTTTC |
| 4CL4 | Peaxi162Scf00610g00346.1 | GAAAGATGAGGTTGCAGGAGAAGTTC | CAGACGGAGATTTTGGGATTGCTTC |
| CA4H | Peaxi162Scf00390g00225.1 | GCCATTGATCACATTCTTGAAGCTCA | CCACCTTTCTCTGTAGTGTCGATCTT |
| C3H1 | Peaxi162Scf00220g00211.1 | TTGAAGGAGTGATGGATGAACAAGGA | AGGTCATATTTCTGTTGGAGGGTCAA |
| C3H2 | Peaxi162Scf00975g00122.1 | AGAACACACTCTTGCTAGGAAGGAA | GAGGCAATCTAGGAGTAGGAACAACT |
| COMT | Peaxi162Scf00912g00111.1 | GATTGGAGTGATGAACATTGCCTGAA | GCTTCAAAGTCCTTCTCAGTCCTTTC |
| CCoAOMT1 | Peaxi162Scf00450g00032.1 | CCTGAACCCATGAAAGAGCTAAGAGA | GCTAGTATCTTGCCATCATCAGGAAG |
| CCoAOMT2 | Peaxi162Scf00016g02023.1 | TGCTTTGCCTGTTCTTGATCTAATGG | AATTTCAATTCTTGGATCAGCAGCCA |
| CCoAOMT3 | Peaxi162Scf00316g00055.1 | GAGGAGTGATAGGCTATGACAACACA | AAAATAAACTAGCTGAGACGCCTGC |
| HCT | Peaxi162Scf00045g02037.1 | CACCAACCCCTTTTACGGATAACAAA | AGTTCCAAACAACAATCTCCCCATTC |
| CAD1 | Peaxi162Scf00016g02329.1 | GGAGTTATCAACACACCCTTGCAATT | CTTCCAGCAACATCGATCACAAATCT |
| CAD2 | Peaxi162Scf00152g00245.1 | TTTGGATTGACGCAGAGTGGATTAAG | ATGTAGTCAAATGAATCAGCAGCCTC |
| CCR1 | Peaxi162Scf00332g00433.1 | GTATGTGCATGTGAAGGATGTAGCTC | TCCAAACCCAAGTCCTTTAGCTTTTG |
| CCR2 | Peaxi162Scf00207g00444.1 | CAGTGCAGGCATATGTGGATGTTAAA | TGTTTCACTGGGGTAAATTCAAGACC |
| F5H1 | Peaxi162Scf00083g01311.1 | TATTAATTCATGGGCCATTGGACGTG | AAGTAAAACAATGAAGAAGGTGGGCC |
| F5H2 | Peaxi162Scf00083g00156.1 | TTATTAATTCATGGGCCATTGGACGC | AAGTAAAACAATGAAGAAGGTGGGCC |

### Table S4 – Relative expression values (vs. GAPDH and actin)

| | AM(mut)/AM(wt) | | | c(mut)/c(wt) | | |
|---|---|---|---|---|---|---|
| | *vpy1** | *vpy2* | *vpy3* | *vpy1** | *vpy2* | *vpy3* |
| PAL-1 | **1.95** | **4.86** | **5.73** | 1.27 | 1.29 | 0.94 |
| PAL-2 | **2.01** | **3.88** | **5.85** | 1.28 | 1.67 | 1.52 |
| PAL-3 | **3.62** | **6.25** | **4.01** | **2.68** | **1.68** | 1.28 |
| 4CL-1 | **5.49** | **4.11** | **6.16** | 0.82 | 1.60 | 1.14 |
| 4CL-2 | **5.21** | **3.37** | **5.39** | 0.81 | 1.52 | 1.19 |
| 4CL-3 | **15.85** | **18.59** | **30.85** | 0.35 | 1.05 | 0.79 |
| 4CL-4 | **6.76** | **21.71** | **16.07** | 1.07 | **0.49** | 0.98 |
| C4H | **4.36** | **5.67** | **3.87** | 0.90 | 0.98 | 1.01 |
| C3H-1 | **5.00** | **5.63** | **6.24** | **0.51** | 0.77 | 0.62 |
| C3H-2 | **3.41** | **3.16** | **5.10** | 0.78 | **1.90** | 1.60 |
| COMT | **3.26** | **7.00** | **8.19** | 0.61 | 1.58 | **1.39** |
| CCoAOMT-1 | **6.82** | **4.39** | **4.24** | 0.59 | **1.88** | 1.52 |
| CCoAOMT-2 | **3.06** | **4.89** | **7.20** | 1.15 | **1.80** | **1.53** |
| HCT | **2.42** | **5.84** | **12.15** | **2.70** | 0.91 | 1.45 |
| CAD1 | **5.59** | **6.58** | **6.22** | 1.47 | 0.96 | 1.08 |
| CAD2 | **8.20** | **15.16** | **26.65** | 0.74 | 0.86 | 0.89 |
| CCR1 | **6.18** | **21.26** | **30.11** | 0.83 | 0.91 | 1.03 |
| CCR2 | **6.18** | **6.64** | **5.80** | 1.06 | 0.88 | 1.17 |
| F5H1 | **5.46** | **16.89** | **23.56** | 0.97 | 0.61 | 0.96 |
| F5H2 | **4.31** | **4.78** | **14.36** | 1.01 | 1.33 | 1.18 |

Table S5 – Relative expression values (vs. GAPDH and actin)

| | non-AM | | AM | |
|---|---|---|---|---|
| | wt | vap-1 | wt | vap-1 |
| PAL-1 | 0.6007 | 0.7640 | 0.6175 | 1.5287 |
| PAL-2 | 1.5103 | 1.9321 | 3.8464 | 9.8658 |
| PAL-3 | 0.1892 | 0.5070 | 0.2403 | 2.3293 |
| 4CL-1 | 0.3082 | 0.2539 | 0.8830 | 3.9912 |
| 4CL-2 | 0.1034 | 0.0832 | 0.1869 | 0.7840 |
| 4CL-3 | 0.0219 | 0.0076 | 0.0207 | 0.1139 |
| 4CL-4 | 0.0364 | 0.0392 | 0.0296 | 0.2151 |
| C4H | 0.5536 | 0.4979 | 0.5332 | 2.0932 |
| C3H-1 | 0.1086 | 0.0550 | 0.1653 | 0.4183 |
| C3H-2 | 0.0027 | 0.0021 | 0.0049 | 0.0130 |
| COMT | 0.1948 | 0.1195 | 0.2101 | 0.4202 |
| CCoAOMT-1 | 0.3435 | 0.2032 | 0.3500 | 1.4112 |
| CCoAOMT-2 | 0.5085 | 0.5852 | 0.6027 | 2.1241 |
| HCT | 0.0008 | 0.0023 | 0.0020 | 0.0131 |
| CAD1 | 0.1851 | 0.2713 | 0.2275 | 1.8643 |
| CAD2 | 0.0598 | 0.0443 | 0.0629 | 0.3819 |
| CCR1 | 0.0685 | 0.0571 | 0.0800 | 0.4118 |
| CCR2 | 0.1595 | 0.1685 | 0.1283 | 0.8371 |
| F5H1 | 0.0358 | 0.0348 | 0.0670 | 0.3564 |
| F5H2 | 0.0867 | 0.0879 | 0.0884 | 0.3858 |

### Table S6 – Relative expression values (vs. GAPDH and actin)

| | non-AM | | | AM | | |
|---|---|---|---|---|---|---|
| | wt | vap-2 | vap-3 | wt | vap-2 | vap-3 |
| PAL-1 | 0.3686 | 0.4755 | 0.3478 | 0.4377 | 2.7429 | 2.3662 |
| PAL-2 | 1.5171 | 2.5267 | 2.3039 | 3.1366 | 20.2450 | 27.8622 |
| PAL-3 | 0.2007 | 0.3372 | 0.2560 | 0.2695 | 2.8288 | 1.3787 |
| 4CL-1 | 0.3710 | 0.5918 | 0.4220 | 0.6853 | 4.4945 | 4.8053 |
| 4CL-2 | 0.0954 | 0.1446 | 0.1135 | 0.1340 | 0.6845 | 0.8596 |
| 4CL-3 | 0.0155 | 0.0163 | 0.0122 | 0.0168 | 0.3290 | 0.4086 |
| 4CL-4 | 0.0280 | 0.0137 | 0.0273 | 0.0202 | 0.2137 | 0.3166 |
| C4H | 0.9613 | 0.9418 | 0.9683 | 1.0128 | 5.6272 | 3.9472 |
| C3H-1 | 0.2986 | 0.2313 | 0.1861 | 0.3585 | 1.5622 | 1.3954 |
| C3H-2 | 0.0091 | 0.0172 | 0.0145 | 0.0094 | 0.0561 | 0.0764 |
| COMT | 0.2357 | 0.3723 | 0.3283 | 0.4523 | 5.0024 | 5.1580 |
| CCoAOMT-1 | 0.2006 | 0.3764 | 0.3051 | 0.3083 | 2.5374 | 1.9880 |
| CCoAOMT-2 | 1.0292 | 1.8533 | 1.5731 | 1.2339 | 10.8633 | 13.5775 |
| HCT | 0.0012 | 0.0011 | 0.0018 | 0.0042 | 0.0224 | 0.0742 |
| CAD1 | 0.0933 | 0.0896 | 0.1011 | 0.2104 | 1.3297 | 1.4182 |
| CAD2 | 0.1732 | 0.1498 | 0.1549 | 0.1914 | 2.5099 | 4.5622 |
| CCR1 | 0.0298 | 0.0271 | 0.0306 | 0.0445 | 0.8588 | 1.3745 |
| CCR2 | 0.0981 | 0.0863 | 0.1144 | 0.1196 | 0.6984 | 0.8097 |
| F5H1 | 0.0246 | 0.0149 | 0.0235 | 0.0260 | 0.2662 | 0.5845 |
| F5H2 | 0.0299 | 0.0399 | 0.0354 | 0.0312 | 0.1988 | 0.5296 |

### Table S7 – qPCR primers for PR genes

| | Gene ID | Forward primer | Reverse primer |
|---|---|---|---|
| PR2a | Peaxi162Scf00032g10002.1 | TTTGATGCCTTTTTGGATTCTATGTA | TCTCACTCTCGTCTCCTTTCTTATCA |
| PR2b | Peaxi162Scf00032g09002.1 | CTGATGTCCCACTATCTTATGCACTT | TATTTTGCAGTTTGATTGGGATAGAA |
| PR2c | Peaxi162Scf00228g00020.1 | AGTTATGGTAGGACTTCCCAATTCAG | GAGTTGCCAATCAAACTCATGTCTAC |
| PR3 | Peaxi162Scf01261g00014.1 | TATGACTTAGCTGGGAAAGCTATTGA | GTAATGACACCATAACCTGGTACACG |
| PR4b | Peaxi162Scf00016g31028.1 | AAATACTAGGACAAGGGCTCAGGTAA | ATTGTCAACTACAGAAAGCAGAGGAA |
| PR4c | Peaxi162Scf00282g05024.1 | CTGTGGTAGATGCTTGAGGAATAAGA | ATAGTTGACAATAAGGTGGCGTTGTT |
| PR4d | Peaxi162Scf00282g05022.1 | TGAATGCTGTTAGTGCTTATTGTTCA | ATTGTCACCACAATTAACAAACTGGT |
| PR5a | Peaxi162Scf00714g01008.1 | CACTTAGGGTACCTGGAGGATGTAAT | AGAGCCATAAGGACAGAAGACAACTT |
| PR6 | Peaxi162Scf00714g02017.1 | TCTTGCATCATTTACTCAACATCTCA | GTCACTGGAGAACCATTCAGTACAGT |
| PR7 | Peaxi162Scf00620g03014.1 | TCCACAGACATATACCAGAACTGTGA | CAAACACTACAGCAATTGGACTTCTT |
| PR9 | Peaxi162Scf00192g00028.1 | AAACACTACGTGGAATATGTCCTCAA | ATATTGCCCATTTTAATCATGGAGTT |
| PR14 | Peaxi162Scf01238g02008.1 | GGTTTAGCTGGTTGTCTTCCTTATTT | GTAAGGAATGTTAACACCACAAGCAG |
| PR17 | Peaxi162Scf00016g24022.1 | AGGAGAGAGATCACTGGCGTATTATT | GTTGCAGTAATCTAGAAATCGAGCAG |
| Actin7 | Peaxi162Scf00025g02112.1 | GAGGTTCCGTTGCCCAGA | CCCGCAGCTTCCATTCC |
| GAPDH | Peaxi162Scf00763g00325.1 | GGAATCAACGGTTTTGGAAGAATTGGGCG | GGCCGTGGACACTGTCATACTTGAACA |

#### Table S8

| | AM (mut)/AM(wt) | | | c(mut)/c(wt) | | |
|---|---|---|---|---|---|---|
| | *vpy-1** | *vpy-2* | *vpy-3* | *vpy-1** | *vpy-2* | *vpy-3* |
| PR2a | **14.47** | **12.05** | **5.22** | 0.87 | 0.66 | 2.27 |
| PR2b | **4.18** | **12.30** | **17.38** | 2.73 | 0.57 | 1.66 |
| PR2c | 1.26 | 0.87 | 0.70 | 0.95 | 1.16 | 0.61 |
| PR3 | **5.63** | **3.61** | **2.24** | 0.53 | 1.31 | 1.65 |
| PR4b | **4.72** | **7.66** | **2.39** | 1.22 | 0.60 | 3.05 |
| PR4c | **9.42** | **9.75** | **12.53** | 0.92 | 1.04 | 1.33 |
| PR4d | **0.43** | 0.49 | 0.29 | **9.78** | **2.14** | **4.23** |
| PR5a | 0.17 | 0.29 | 0.33 | **2.40** | 0.98 | **2.74** |
| PR6a | 3.40 | 2.55 | 0.12 | **6.71** | **15.97** | **15.59** |
| PR7 | **6.29** | **6.45** | **1.35** | 0.29 | 1.01 | **1.48** |
| PR9 | **3.52** | **1.22** | **4.97** | 0.83 | 1.33 | 1.36 |
| PR14 | **5.90** | 7.15 | **6.56** | 0.84 | 0.64 | 1.27 |
| PR17 | **6.67** | **6.18** | **1.76** | 1.57 | 1.59 | **2.25** |

#### Table S9

| | 1h | | 4h | |
|---|---|---|---|---|
| | Chit | Pen | Chit | Pen |
| PR2a | 1.38 | 2.93 | 0.61 | 0.82 |
| PR2b | 1.52 | 0.88 | 0.82 | 0.50 |
| PR2c | 1.38 | 1.35 | **2.61** | **3.37** |
| PR3 | 1.91 | 1.33 | 0.93 | **1.78** |
| PR4b | 0.89 | 1.64 | 1.15 | **2.44** |
| PR4c | 2.04 | 0.88 | 0.97 | 0.92 |
| PR4d | 1.61 | 1.61 | **5.45** | **5.02** |
| PR5a | 1.45 | 2.21 | 1.54 | **3.26** |
| PR6a | 0.73 | 0.61 | 0.92 | **2.84** |
| PR7 | 2.99 | 1.02 | 0.94 | 1.34 |
| PR9 | 2.16 | 1.22 | 1.27 | 1.17 |
| PR14 | 1.33 | 0.87 | 1.02 | 1.07 |
| PR17 | 1.55 | 0.64 | 0.99 | **1.96** |

Yellow: induction>2, orange, induction>4 ; bold: $p<0.05$

Table S10 – Relative expression values (vs. GAPDH and actin)

| | non-AM | | | AM | | |
|---|---|---|---|---|---|---|
| | *wt* | *vpy-2* | *vpy-3* | *wt* | *vpy-2* | *vpy-3* |
| PR2a | 0.0008 | 0.0005 | 0.0018 | 0.0020 | 0.0158 | 0.0235 |
| PR2b | 0.0003 | 0.0001 | 0.0004 | 0.0004 | 0.0025 | 0.0103 |
| PR2c | 0.5561 | 0.6429 | 0.3387 | 0.7008 | 0.7030 | 0.2983 |
| PR3 | 0.0119 | 0.0156 | 0.0196 | 0.0165 | 0.0782 | 0.0609 |
| PR4b | 0.3502 | 0.2093 | 1.0678 | 1.9080 | 8.7328 | 13.8989 |
| PR4c | 0.0032 | 0.0033 | 0.0042 | 0.0035 | 0.0355 | 0.0580 |
| PR4d | 0.1799 | 0.3843 | 0.7605 | 0.9921 | 1.0316 | 1.2303 |
| PR5a | 3.5041 | 3.4324 | 9.5838 | 12.1965 | 3.4931 | 10.8562 |
| PR6a | 0.0011 | 0.0183 | 0.0179 | 0.0152 | 0.6182 | 0.0281 |
| PR7 | 0.0050 | 0.0050 | 0.0074 | 0.0260 | 0.1687 | 0.0522 |
| PR9 | 0.0015 | 0.0020 | 0.0020 | 0.0079 | 0.0128 | 0.0532 |
| PR14 | 0.0038 | 0.0024 | 0.0048 | 0.0114 | 0.0527 | 0.0949 |
| PR17 | 0.1266 | 0.2012 | 0.2844 | 0.1771 | 1.7394 | 0.7017 |

Table S11 – Relative expression values (vs. GAPDH and actin)

| | non-AM | | AM | |
|---|---|---|---|---|
| | *wt* | *vpy-1* | *wt* | *vpy-1* |
| PR2a | 0.0006 | 0.0005 | 0.0008 | 0.0098 |
| PR2b | 0.0002 | 0.0005 | 0.0003 | 0.0039 |
| PR2c | 0.2096 | 0.1985 | 0.3112 | 0.3707 |
| PR3 | 0.0169 | 0.0090 | 0.0197 | 0.0593 |
| PR4b | 0.7377 | 0.8976 | 1.5068 | 8.6525 |
| PR4c | 0.0012 | 0.0011 | 0.0011 | 0.0094 |
| PR4d | 0.2050 | 2.0049 | 1.1415 | 4.7825 |
| PR5a | 2.0615 | 4.9515 | 13.7566 | 5.6603 |
| PR6a | 0.0222 | 0.1489 | 0.0399 | 0.9097 |
| PR7 | 0.0099 | 0.0028 | 0.0421 | 0.0762 |
| PR9 | 0.0013 | 0.0011 | 0.0092 | 0.0270 |
| PR14 | 0.0024 | 0.0020 | 0.0060 | 0.0298 |
| PR17 | 0.2138 | 0.3355 | 0.2889 | 3.0270 |

Table S12 – Relative expression values (vs. GAPDH and actin)

| | 1h | | | 4h | | |
|---|---|---|---|---|---|---|
| | **cont** | **CHIT** | **PEN** | **cont** | **CHIT** | **PEN** |
| **PR2a** | 0.004 | 0.006 | 0.013 | 0.016 | 0.010 | 0.013 |
| **PR2b** | 0.001 | 0.002 | 0.001 | 0.004 | 0.003 | 0.002 |
| **PR2c** | 1.825 | 2.514 | 2.467 | 6.542 | 17.080 | 22.041 |
| **PR3** | 0.006 | 0.011 | 0.007 | 0.009 | 0.008 | 0.015 |
| **PR4b** | 1.126 | 1.006 | 1.848 | 1.971 | 2.269 | 4.800 |
| **PR4c** | 0.002 | 0.004 | 0.002 | 0.007 | 0.006 | 0.006 |
| **PR4d** | 0.679 | 1.091 | 1.091 | 1.709 | 9.311 | 8.569 |
| **PR5a** | 5.513 | 7.994 | 12.161 | 19.895 | 30.722 | 64.844 |
| **PR6a** | 0.579 | 0.423 | 0.356 | 1.023 | 0.937 | 2.906 |
| **PR7** | 0.028 | 0.083 | 0.028 | 0.062 | 0.058 | 0.083 |
| **PR9** | 0.005 | 0.011 | 0.006 | 0.012 | 0.016 | 0.015 |
| **PR14** | 0.041 | 0.055 | 0.036 | 0.050 | 0.051 | 0.053 |
| **PR17** | 0.173 | 0.269 | 0.110 | 0.204 | 0.203 | 0.402 |
